# Network science inspires novel tree shape statistics

**DOI:** 10.1101/608646

**Authors:** Leonid Chindelevitch, Maryam Hayati, Art F. Y. Poon, Caroline Colijn

## Abstract

The shape of phylogenetic trees can be used to gain evolutionary insights. A tree’s shape specifies the connectivity of a tree, while its branch lengths reflect either the time or genetic distance between branching events; well-known measures of tree shape include the Colless and Sackin imbalance, which describe the asymmetry of a tree. In other contexts, network science has become an important paradigm for describing structural features of networks and using them to understand complex systems, ranging from protein interactions to social systems. Network science is thus a potential source of many novel ways to characterize tree shape, as trees are also networks. Here, we tailor tools from network science, including diameter, average path length, and betweenness, closeness, and eigenvector centrality, to summarize phylogenetic tree shapes. We thereby propose tree shape summaries that are complementary to both asymmetry and the frequencies of small configurations. These new statistics can be computed in linear time and scale well to describe the shapes of large trees. We apply these statistics, alongside some conventional tree statistics, to phylogenetic trees from three very different viruses (HIV, dengue fever and measles), from the same virus in different epidemiological scenarios (influenza A and HIV) and from simulation models known to produce trees with different shapes. Using mutual information and supervised learning algorithms, we find that the statistics adapted from network science perform as well as or better than conventional statistics. We describe their distributions and prove some basic results about their extreme values in a tree. We conclude that network science-based tree shape summaries are a promising addition to the toolkit of tree shape features. All our shape summaries, as well as functions to select the most discriminating ones for two sets of trees, are freely available as an R package at http://github.com/Leonardini/treeCentrality.

## 2 Introduction

Molecular data describing the evolution, variation and diversity of organisms over time is more widely available than ever before due to rapid improvements in sequencing technology. Using these data to infer the underlying evolutionary process is a key ongoing challenge in many areas of biology. In particular, in infectious disease, it is crucial to understand pathogen adaptation: despite improvements in sanitation and vaccination and the development of antibiotics, infectious pathogens continue to emerge from zoonotic infections and to adapt to human immune responses, vaccines, and antimicrobials. Next-generation sequencing has afforded unprecedented opportunities to generate pathogen genome sequences in a highly scalable manner, and theoretical tools have been developed to interrogate these data, largely through reconstructed phylogenetic trees.

There has been considerable interest over the years in comparing the shapes of phylogenetic trees in order to understand evolutionary processes [64, 63, 24, 58, 31, 47, 6, 71]. A tree’s shape specifies its connectivity structure. The lengths of its branches typically reflect either the time or genetic distance between branching events. Following the observation that reconstructed evolutionary trees are more asymmetric than random models predict [2], there have been efforts to summarise tree asymmetry in trees reconstructed from data and relate it to predicted asymmetry in evolutionary and ecological models [31, 19, 3, 55, 66, 47, 6, 58, 38]. There is also interest in establishing whether taxa from two phylogenies might correspond to each other, for example in the context of parasites and hosts or fossils of different origins [22], and in comparing simulated trees with trees from data in epidemiology, for example using Approximate Bayesian Computation [56, 62]. These applications require quantitative tools to compare phylogenetic trees with different taxa, and they require summary features that are informative of the evolution or epidemiology being studied.

Summaries of tree topology have often focused on either asymmetry, or the frequency of various configurations such as cherries or ladders [60]. Well-known measures of asymmetry include the Colless and Sackin imbalance [12, 61]. Asymmetry measures tend to be correlated with each other, and do not fully capture the shape of a tree [22, 40], leading to an interest in exploring other statistics, comparisons tools and metrics for this task [27, 57, 36, 22, 11, 62]. Some metric approaches directly find a distance (or similarity) measure between unlabelled phylogenies; others seek an optimal labelling for two unlabelled trees and use metrics for labelled trees, but this is not feasible for large trees. Metric approaches also do not lend themselves to interpretable descriptions of trees that can easily be connected with generative models of evolution or epidemiology.

In one particularly important application of tree shape statistics, namely, Approximate Bayesian Computation, one seeks simulation parameters that produce trees that are similar enough to data-derived trees [14]. This leaves a need for new statistics to quickly provide a good summary of tree topology. Matsen used optimization over binary recursive tree shape statistics to find statistics that distinguish between sets of trees [41]. This set the stage for broadening the set of quantities used to describe trees, and gives rise to natural questions like why a particular recursive statistic separates two groups, whether it only happens to separate those specific trees or acts as a good way to distinguish between trees that are meaningfully similar to those in the two groups. The binary recursive class may furthermore exclude features that capture global information about a tree, such as features derived from its eigenvalues when viewed as a graph, known as spectral features. As the size of datasets and range of applications increases, it is reasonable to assume that inference will be improved by expanding upon the tools available to summarize tree topologies [41, 62]. In applications like Approximate Bayesian Computation, as well as other related attempts at summarizing tree shapes, one does not necessarily seek to relate summaries to evolutionary mechanisms. Rather, the key objective is to distinguish between trees in different categories or scenarios in an efficient way by computing simple statistics.

Network science has become an important paradigm for describing structural (topological) features of networks and using them to understand complex systems, ranging from protein interactions to social systems [35, 49]. Network science is thus a potential source of many novel ways to characterize tree topology. A phylogeny can be interpreted as a simple type of network or graph – specifically, a connected acyclic undirected graph. Thinking of a tree as a graph can lead to a matrix representation (such as the adjacency matrix) in which both rows and columns correspond to its internal and terminal nodes. This makes available the large assortment of techniques for studying the properties of graphs based on their matrix representation [25]. The *spectra* of graph matrices have recently been used to compare tree topologies [36]. However, the use of classical spectra is not likely to uniquely define trees [43, 42], the computation does not scale well with tree size, and describing the relationship between evolutionary or epidemiological parameters and graph spectra may be extremely difficult due to the algebraic rather than combinatorial nature of the latter.

Network science offers many other network features that can be adapted to describe shape. Here, we tailor tools from network science to summarize phylogenetic tree topologies. We thereby develop tree topology summaries that are complementary to both asymmetry and the frequencies of small configurations. These new statistics are fast to compute and will scale well to describe the topologies of large trees. They can additionally be easily adapted to take branch lengths into account. We illustrate how these statistics vary, alongside some conventional tree statistics, over three very different viruses (HIV, dengue fever and measles), and over the same virus in different epidemiological scenarios (influenza A over a 2-year period in the USA, over a 5-year period globally, and over a longer period globally; HIV in three epidemiological contexts, namely a concentrated epidemic in men who have sex with men, a generalised African HIV epidemic in a village setting and HIV-1 from a national-level survey (see Methods)). We also use the statistics to compare simulated trees from different models. Finally, we use classification with and without the network-based statistics to distinguish trees in different settings. We find that the network-science statistics improve the classification performance and are consistently assigned a high importance measure in classification algorithms.

## 3 Methods

### 3.1 Definitions related to graphs

In this subsection we introduce some definitions from graph theory, used below. They are based on the textbooks by Bollobás [7] as well as Godsil and Royle [20].

A *network* or *graph G* consists of a finite set of *nodes V* and a finite set of *edges E* whose elements are unordered pairs of distinct nodes. The edge {*u, v*}, denoted *uv* for simplicity, *joins* nodes *u* and *v*, and if *uv* ∈ *E*, the nodes *u* and *v* are *adjacent*, which we denote by *u* ~ *v*.

A graph is *weighted* if there is a function *w* : *E* → ℝ that assigns a *weight w*(*uv*) to each edge *uv* ∈ *E*. The weight function extends to all of *V* × *V* by setting *w*(*uv*) = 0 if *u* = *v* or *uv* ∉ *E*.

A *path P* in a graph is a sequence of nodes *v*_0_, *v*_1_, …, *v_k_* such that *v_i_v*_*i*+1_ ∈ *E* for each 0 ≤ *i* ≤ *k* − 1. The nodes *v*_0_ and *v_k_* are the *endvertices*. In the unweighted case, *k* is the *length* of *P*. In the weighted case, the weighted length of the path *P* is measured by the sum of the weights of the edges it contains, i.e. 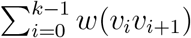.

The path *P* corresponding to the sequence *v*_0_, *v*_1_, …, *v_k_* is a *simple path* if *v_i_* ≠ *v_j_* for any 0 ≤ *i* < *j* ≤ *k*. If the endvertices are identical, i.e. *v*_0_ = *v_k_*, then *P* is a *cycle*; if the endvertices are the only identical nodes on the cycle, it is a *simple cycle*. A simple cycle of length 2 is just an edge traversed twice; a graph with no simple cycle of length greater than 2 is called *acyclic*.

In both the unweighted and the weighted cases, the minimum (weighted) length of a path with endvertices *u* and *v* is called the (weighted) *distance* between *u* and *v* and denoted *d*(*u, v*). The (weighted) *diameter* of the graph is the largest distance between two distinct nodes.

Being the endvertices of some path of *G* defines an equivalence relation on the set of nodes, and its equivalence classes are called the *connected components* (or simply *components*) of *G*. A graph is *connected* if it contains only one component. A node whose deletion increases the number of components is a *cutvertex* ; an edge whose deletion increases this number is a *bridge*.

The *neighborhood* Γ(*u*) of a node *u* ∈ *G* is the set of nodes adjacent to it in *G*. The *degree* of *u* is *d*(*u*) = |Γ(*u*)|. The *degree sequence* of *G* is the vector **d** = (*d*(*v*_1_) … *d*(*v_n_*)) containing the degree of all its nodes in some order. In a weighted graph, the *weighted degree* of *u* is *d_w_*(*u*) = Σ_*v*_ *w*(*uv*), and the *weighted degree sequence* is the vector **d_w_** = (*d_w_*(*v*_1_) … *d_w_*(*v_n_*)). The *density* of *G* is the ratio *ρ* ≔ |*E*| / |*V*|; if *G* is unweighted, *ρ* equals half the average degree.

The *clustering coefficient* of a node *u* in a graph is defined as 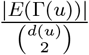, where *E*(Γ(*u*)) is the set of pairs *v, v*′ ∈ Γ(*u*) such that *vv*′ ∈ *E*. This coefficient measures the fraction of mutually adjacent nodes among the nodes adjacent to *u*, and varies from 0 to 1. The *assortativity coefficient* of *G* is the Pearson correlation coefficient of degree between pairs of adjacent nodes.

Given a numbering of the nodes, *V* = {1, 2, …, *n*}, a graph can be represented by its *adjacency matrix A* ∈ ℝ^*n*×*n*^, where *A_ij_* = 1 if *ij* ∈ *E* and *A_ij_* = 0 otherwise. If the graph is weighted, its *weighted adjacency matrix* has entries *A_ij_* = *w*(*ij*).

A graph can also be represented by its *Laplacian matrix L* ∈ ℝ^*n*×*n*^, defined by *L_ii_* = *d*(*i*) for all *i*, *L_ij_* = −1 if *ij* ∈ *E* and *L_ij_* = 0 otherwise. The *weighted Laplacian matrix* has entries *L_ii_* = *d_w_*(*i*) for all *i* and *L_ij_* = −*w*(*ij*) if *i* ≠ *j*. Thus, in both cases we have *L* = *D* − *A*, where *D* is the diagonal matrix with the (weighted) degree sequence on the diagonal.

The *normalized Laplacian* [9] is a variant of the Laplacian which is normalized by the degree of each node, and is defined as 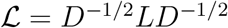. Its entries are 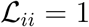 for all *i* and 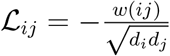 for *i* ≠ *j* in the weighted case or 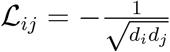 in the regular case.

In addition, a graph can also be represented by its *distance matrix Q* ∈ ℝ^*n*×*n*^, with *Q_ij_* = *d*(*i, j*), the (weighted or unweighted) graph distance between *i* and *j*, and in particular, *Q_ii_* = 0. As proposed in [36], it is also possible to apply the (regular or normalized) Laplacian transformation to the distance matrix (by treating the distance matrix as a weighted adjacency matrix) to obtain alternative matrix representations of a graph.

Widely studied graph properties fall into two broad classes. The first are *shape properties*, concerned with various connectivity measures of the graph, such as the numbers of distinct paths connecting pairs of nodes and the average or longest distance between a pair of nodes. Some of these properties do not depend on the edge weights of the graph, in which case we refer to them as *topology properties*; we use the term *shape property* as a broad term that includes both topology properties and properties that take edge weights into account. The second are *spectral properties*, concerned with the eigenvalues (and occasionally eigenvectors) of matrices representing the graph, such as the adjacency, Laplacian, and normalized Laplacian.

### 3.2 Definitions related to trees

A tree is a connected acyclic graph. It can be shown that a tree with *n* nodes has *n* − 1 edges, and that there is a unique simple path between any two nodes in a tree. The nodes of degree one are called *tips* (or *leaves*); the remaining nodes are called *internal*. The weight *w*(*e*) of an edge *e* of a tree is also called its *branch length*; we assume that a tree has all positive branch lengths.

A tree *T* is *rooted* if it contains a distinguished node *r*, called the *root*. If *v* is any node other than the root *r* in a rooted tree, the node *u* immediately preceding *v* on the unique simple path from *v* to *r* is called the *parent* of *v* in *T*, and any node *z* on this unique simple path is called an *ancestor* of *v* in *T*. In particular, the root *r* is an ancestor of every node in *T* other than itself. The number of edges on the unique shortest path from a node *v* to the root *r* is called its *depth*.

Conversely, in this situation, *v* is called a *child* of *u* and a *descendant* of *z*. A rooted tree *T* is *binary* if every internal node has exactly two children. In a rooted tree *T*, the set of all descendants of a node *u* including *u* itself forms a *clade*; we say that *u subtends* this clade. The *subtree* rooted at *u*, denoted *T_u_*, is the clade that *u* subtends together with all the edges of *T* connecting its elements. If a node *u* has two children *v* and *w*, we may arbitrarily refer to them as the left and right child of *u*, and call *T_v_* and *T_w_* the left and right subtree of *u*, respectively.

A tree *T* with root node *r* can be *rerooted* at an internal node *s* ≠ *r*, by simply designating *s* rather than *r* as the root. This changes only the ancestor-descendant relationships, but not the topology of the tree. Note that if *T* is a binary tree with root *r*, *T* rerooted at *s* ≠ *r* will not be binary because *s* will have three children while *r* will have only one child after rerooting. However, this situation is often resolved by specifying a branching order, i.e. creating a new node *t* that is the parent of two children of the new root *s*, and adding an *s* − *t* edge of length 0. An analogous procedure, called “multichotomy resolution”, may be iteratively applied to any other internal nodes with a degree greater than 2 to turn any non-binary tree into a binary tree.

A molecular *phylogeny*, or *phylogenetic tree*, is a binary rooted tree whose tips correspond to genetic or genomic sequences, and whose internal nodes represent their inferred common ancestors. A phylogeny therefore represents the ancestral relationships among a set of genomes. The shape of a phylogeny tells the story of an evolutionary history going back through time to the most recent common ancestor at the root of the tree (see Figure 1 (a)).

**Figure 1:**
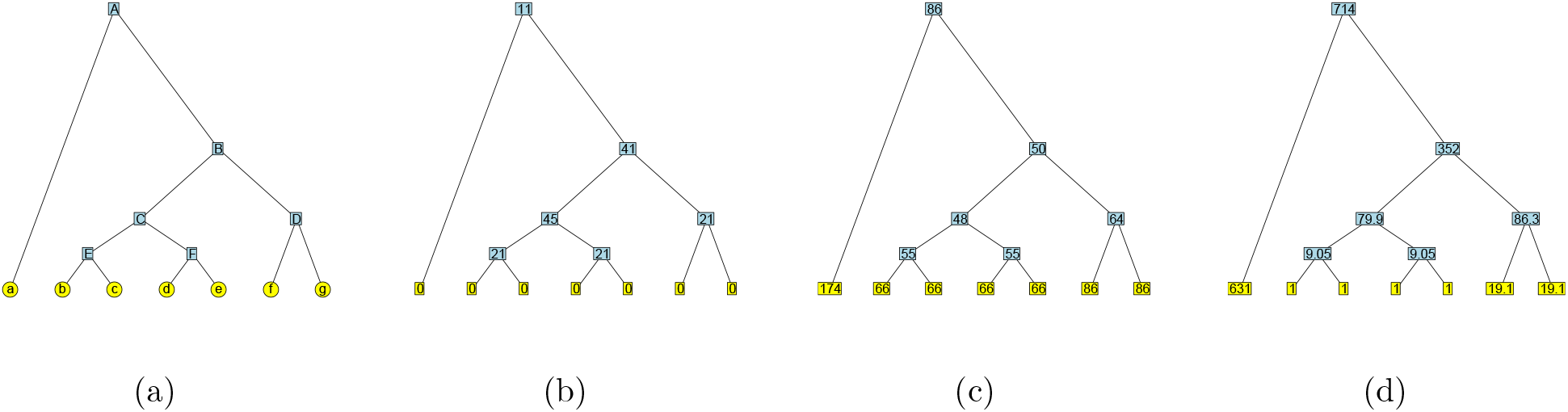
(a) An example tree. In the epidemiological context, tips (*a* – *g*) would correspond to pathogen sequences and internal nodes (*A* – *F*) to their inferred common ancestors. Node *D* subtends a “cherry” configuration, and node *C* subtends two cherries (a “double cherry”). The heights of the internal nodes are 1 (*E, F*), 2 (*C, D*), 4 (*B*) and 8 (*A*), so the diameter is 16 and the Wiener index is 484, for a mean path length of 6.21. (b) Same tree with betweenness centrality values at each node (note that branch lengths do not change them). The tree has betwenness centrality 45. (c) Same tree with farness (reciprocal of closeness centrality) values at each node. The tree has closeness centrality 1/48. (d) Same tree with eigenvector centrality values (scaled to have a minimum of 1) at each node, rounded to 3 significant figures. Here, the leading eigenvalue is *λ* = 9.05. By our definition, the tree has eigenvector centrality 714/1023 = 0.698 (here, 1023^2^ is the sum of the squared values).

### 3.3 Data and simulations

#### HIV/Dengue/Measles

We obtained Newick tree strings corresponding to phylogenies inferred from human and zoonotic RNA viruses from a previous study. Specifically, we retrieved tree strings reconstructed from genetic sequences of HIV-1 subtype B, Measles virus, and Dengue virus serotype 4. The HIV sequence data (corresponding to the gene encoding the Nef protein) were obtained from the LANL HIV Sequence database [18], through the web site at http://www.hiv.lanl.gov, and screened for recombinants using the SCUEAL algorithm [32]. The remaining virus sequences were obtained from GenBank [4].

Phylogenies were reconstructed from random samples of 100 sequences by maximum likelihood using RAxML [65] under a general time-reversible model of nucleotide substitution and rate variation among sites approximated by the GTRCAT model. HIV-1 subtype B phylogenies were rooted using a subtype D sequence as an outgroup. Dengue virus serotype 4 phylogenies were rooted on an outgroup sequence isolated in the Philippines in 1956. Finally, measles virus phylogenies were rooted using a genotype D6 sequence as the outgroup. The GenBank [4] accession numbers for all outgroups can be found in Table S1 in the **Supplementary Materials**.

#### HIV in three settings

We obtained HIV-1 sequence data from three published studies. The Wolf et al. [70] data set corresponds to samples from a concentrated epidemic of HIV-1 subtype B in populations of predominantly men who have sex with men in Seattle, USA. The Novitsky et al. [53] data set corresponds to samples from a generalized epidemic of HIV-1 subtype C infections in Mochudi, Botswana, a village with an estimated HIV-1 prevalence of about 20% in the adult population. Similarly, the Hunt et al. [28] data set represents samples from a national survey of the generalized epidemic of HIV-1 subtype C in South Africa. Thus, these studies represent a range of geographic scales and epidemiological contexts.

#### Influenza in three settings

We compared the topologies of a single virus (human influenza A) sampled to reflect different epidemiology: (1) samples over a five-year period; (2) global samples over a 12-year period and (3) samples from 2012-2013 from the USA only. We downloaded full-length hemagglutinin (HA) sequences of human H3N2 flu from NCBI and aligned sequences from each group with MAFFT [30]. For each sample, we chose 120 sequences uniformly at random from the alignment, and inferred a tree with these sequences as tips using IQtree [67] with the pll (phylogenetic likelihood library) option [17] and the GTR+G model. Using the date information from NCBI, we rooted the trees using the root to tip (*rtt*) function in the *ape* package [54] in R [59].

#### Simulated tree models

We simulated trees from four random processes: a Yule process (pure birth trees), a “biased” model of Kirkpatric and Slatkin [31] in which speciation rates are unevenly assigned to a node’s descendants with a bias (here 0.3), and two constant rate birth-death processes, with the basic reproduction number (mean of the offspring distribution) equal to 1.5 and 3. We created sets of 100 trees with 100 tips and separately with 300 tips. The `apTreeshape` package was used to simulate the Yule and biased models; tree shapes were converted to phylogenetic trees using the `as.phylo` function. We used the `TreeSim` package for the birth-death models. Because in `sim.bd.taxa` (in `TreeSim`) the simulations are conditioned on having a fixed number of *extant* tips, we created trees with 300 or 600 extant tips and randomly pruned taxa to leave a tree of 100 or 300 tips, modelling partial sampling over time.

As some scenarios can be distinguished simply by comparing the branch lengths of the corresponding trees, we normalized the time scales so that each of our trees has a mean branch length of 1. This ensures that any differences we observe between the summary statistics in different classes are not simply due to scaling. We did not, however, modify the variances of the branch length distribution, as those may contain some of the signal picked up by summary statistics.

In total, there are 5 scenarios in which we compare trees: HIV/Dengue/Measles (HDM), influenza (2-year USA, 5-year global, 12-year global), HIV contexts (labeled WNH after the first author names of the corresponding publications), simulated trees with 100 tips (’Simulated’) and simulated trees with 300 tips (’Simulated300’). Within each set, we performed classification with generalized linear models and random forests, using the tree shape statistics as features. We computed a measure of each feature’s importance for each classification. All reconstructed trees, including accession numbers (in the tip labels), have been deposited to github.com/Leonardini/treeCentralityData.

### 3.4 Shape features

We computed a range of topological and spectral summary features of the viral phylogenies (see Table 1 for the definitions and references for each one). Our focus here is on some of the novel tree topology summary statistics, but we also include a number of standard statistics for comparison. All input trees were binary and rooted, and all branch lengths were non-negative, although many of the trees had zero-length branches. All comparisons involved trees on the same number of tips.

**Table 1:** Summary measures for phylogenetic trees. Here, *n_i_* is the number of nodes at depth *i*.

| Name | Description | Short form | Ref. |
| --- | --- | --- | --- |
| Numbers of small configurations |  |  |  |
| Cherry number | # of nodes with 2 tip children | cherries | [44] |
| Pitchforks | # of nodes with 3 tip descendants | pitchforks | [60] |
| Double cherries | # of nodes with 2 cherry children | doubcherries | new |
| 4-caterpillar | # of caterpillars with 4 tips | fourprong | [60] |
| Clades of size $x$ | # of nodes with $x$ tip descendants | num$x$ | [60] |
| Tree-wide summaries |  |  |  |
| Colless imbalance | Colless imbalance | colless | [12] |
| Sackin imbalance | Mean path length from tip to root | sackin | [61] |
| Maximum height | Max # of steps from the root | maxheight | [10] |
| Maximum width | Max # of nodes at the same depth | maxwidth | [10] |
| Stairs | Proportion of imbalanced subtrees | stairs | [52] |
| Max difference in widths | $\max_i(n_{i+1} - n_i)$ | delW | [10] |
| Node properties from network science |  |  |  |
| Betweenness centrality | # of shortest paths through node | between | [51] |
| Weighted betweenness | as above, but with weighted edges | betweenW | [50] |
| Closeness centrality | total distance to all other nodes | closeness | [51] |
| Weighted closeness | as above, but with weighted edges | closenessW | [50] |
| Eigenvector centrality | value in Perron-Frobenius vector | eigen | [51] |
| Weighted eigenvector | as above, but with weighted edges | eigenW | [50] |
| Summaries from network science |  |  |  |
| Diameter | largest distance between 2 nodes | diameter | [7] |
| Mean pairwise distance | average distance between 2 nodes | meanpath | [45] |
| Spectral properties |  |  |  |
| Min adjacency | min adjacency matrix eigenvalue $> 0$ | minAdj | [20] |
| Max adjacency | max adjacency matrix eigenvalue | maxAdj | [20] |
| Min Laplacian | min Laplacian matrix eigenvalue $> 0$ | minLap | [20] |
| Max Laplacian | max Laplacian matrix eigenvalue | maxLap | [20] |
| Distance Laplacian spectral properties |  |  |  |
| Max eigenvalue | largest eigenvalue in the spectrum | dLapLambdaMax | [36] |
| Max density | location of largest spectral density | dLapDensityMax | [36] |
| Asymmetry | skewness of the spectral density | dLapAsymmetry | [36] |
| Kurtosis | peakedness of the spectral density | dLapKurtosis | [36] |

For the node properties derived from network science, we focus our discussion on the maximum value of each type of centrality a node can have within a tree, but using other derived statistics (the minimum, mean, median or variance) could also have been an option. For the spectral properties, which also produce a value for each node, we focus on the maximum value as well as the minimum strictly positive value. Lastly, for the distance Laplacian spectral properties we use the four derived statistics proposed by Lewitus and Morlon [36].

#### 3.4.1 Tree topology summaries based on network science

Network science, broadly defined as the study of complex networks, has produced a number of tools that have been applied in a variety of contexts [35, 49], including degree sequence, degree assortativity, density, diameter, and a number of node centrality measures. Only some of these apply naturally and informatively to phylogenies. For instance, the degree sequence, the degree centrality, the density and the clustering coefficients do not vary between phylogenetic trees on *n* tips, while their degree assortativity can take on one of two values which are close to one another for large *n* (see section 6.4 in the **Supplementary Materials** for details).

We now discuss those network science-inspired features that are informative for phylogenetic trees.

#### 3.4.2 Diameter, average shortest path and Wiener index

The diameter (the maximum length of a shortest path) is a useful summary statistic. For general trees with *N* nodes, it can be calculated in linear time using a “folklore” dynamic programming algorithm, and its value can vary between 2 for the star and *N* − 1 for the path of length *N*. For phylogenetic trees with *N* = 2*n* 1 nodes, the range is from 2 log(*n*) to *n*. The exact maximum and minimum values, as well as the trees attaining them, and the distribution for *N* = 45 (*n* = 23), are described in Figure S10 in **Supplementary Materials**.

The average shortest path length of a tree may also be informative. Unlike for general graphs, it can be calculated in linear time in a tree with *N* nodes using dynamic programming [45]. The sum of all shortest path lengths between pairs of nodes in a tree is known as the tree’s Wiener index, and equals (2*n* − 1)(*n* − 1) times the average shortest path length. The Wiener index for general trees with *N* nodes is always contained between (*N* − 1)^2^ and 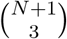, values which are attained by the star and the path, respectively [16]. For phylogenetic trees, the range of the Wiener index is substantially narrower; its distribution for *N* = 45 (*n* = 23) is described in Figure S11 in the **Supplementary Materials**. Note that both the diameter and Wiener index generalize naturally to trees with arbitrary positive branch lengths.

Based on their distributions for small phylogenetic trees, shown in Figure S12 in the **Supplementary Materials** we can think of trees that have a diameter or a Wiener index much smaller than the mean as being similar to complete binary trees, and of those that have a diameter or a Wiener index much larger than the mean as being similar to caterpillars or double caterpillars, respectively. This is not too different from the corresponding situation for imbalance statistics such as Colless or Sackin index. However, the situation changes dramatically when we look at node centrality measures defined in network science. A number of such centrality measures exist that can be defined for each node of a graph. In particular, betweenness centrality, closeness centrality and eigenvector centrality are quantities derived from network science that can be computed in linear time for a tree with *N* nodes and can capture aspects of tree topology not captured by mere asymmetry.

#### 3.4.3 Betweenness centrality

Betweenness centrality associates to each node *v* in a graph the number of pairs *u, w* ∈ *V* − {*v*} such that the shortest *u* − *w* path passes through *v*; in other words,

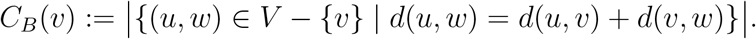

Betweenness centrality can be normalized by the number 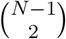 of all pairs *u, w* ∈ *V* − {*v*}, but we choose not to use this normalization here. In a tree *T*, there is a unique shortest path between every pair of nodes, and the shortest *u* − *w* path passes through *v* if and only if *u* and *w* are in clades subtended by different children of *v* when *T* is rerooted at *v*. Hence, the betweenness centrality of an internal node *v* is simply Π_*i*<*j*_*n_i_n_j_*, where *n*_1_, …, *n_k_* are the sizes of the clades subtended by the *k* children of *v* when *T* is rerooted at *v*, and *k* is the degree of *v* in *T*. This implies that the betweenness centrality of all the nodes in a tree can be computed in linear time, a result that, while not surprising, does not seem to have been mentioned in the literature until now, although an algorithm to perform this computation in a distributed fashion has been published [68]. This contrasts with the situation for a general graph, where the best algorithm takes *O*(*NM*) time, where *M* is the number of edges, making it quadratic time for trees [8].

In a phylogenetic tree, the degree is 1 for a tip, 2 for the root and 3 for internal nodes, so the betweenness centrality is respectively 0, *n*_1_*n*_2_ or *n*_0_*n*_1_ + *n*_0_*n*_2_ + *n*_1_*n*_2_ in those cases, where *n*_1_ and *n*_2_ are the sizes of the left and right subtrees of the node and *n*_0_ is the number of nodes outside the subtree rooted at this node. This can easily seen to be maximal when *n*_0_ = *n*_1_ = *n*_2_, a situation that does not occur in every tree, but is only possible in those with *N* ≡ n *n* ≡ 1 mod 3. The betweenness centralities are shown for all the nodes of an example tree in Figure 1 (b). Trees with high maximum betweenness centrality are those which have a node whose left subtree, right subtree, and “outside subtree” (what remains of the tree after removing the subtree rooted at the node), are all close in size - a kind of three-way symmetry, as opposed to the two-way (left-right) symmetry measured by classical statistics. The distribution of this quantity, maximum betweenness centrality, for *N* = 45 (*n* = 23), is described in Figure S13 in the **Supplementary Materials**. We also note that the betweenness centrality values of a tree do not change if the tree has variable branch lengths, since each pair of nodes is connected by a unique shortest path. However, this observation is not true for more complicated graphs.

#### 3.4.4 Closeness centrality

Closeness centrality associates to each node *v* in a graph the inverse of the sum of its distances to all the other nodes in the graph. In other words,

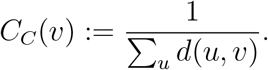

The definition means that closeness centrality is inversely proportional to *farness*, the sum of distances from a node to all the other nodes in the graph; hence, it will be small for centrally located nodes and large for remote ones. While this quantity generally requires at least *O*(*NM*) time to be computed for a graph with *N* nodes and *M* edges [5], this can be reduced to linear time for a tree, an observation that does not seem to have been published previously, although a distributed algorithm for performing this computation has been proposed [69].

Indeed, if we consider an internal node *u* with a child *v*, we have *d*(*v, x*) = *d*(*u, x*) − 1 for every node *x* in the clade of *T* that *v* subtends, and *d*(*v, y*) = *d*(*u, y*) + 1 for every node *y* outside this clade. Hence, by computing the height (distance to the root) of each node and the sizes of the left and right subtrees of each node in a bottom-up traversal of the tree, we can also find the closeness centrality in linear time for all the nodes in the tree. This is illustrated on our example tree in Figure 1 (c). It can also be shown - as we do in Section 6.7 in the **Supplementary Materials** - that the largest value of closeness centrality in a phylogenetic tree is achieved by its unique centroid (the node *v* such that, if the tree is rerooted at it, no child of *v* subtends a clade with more than half of all the nodes), while the smallest value is achieved by one of the tips (in fact, this result also holds for trees with arbitrary positive branch lengths). The distribution of this quantity, maximum closeness centrality, for *N* = 45 (*n* = 23), is described in Figure S14 in the **Supplementary Materials**.

#### 3.4.5 Eigenvector centrality

Eigenvector centrality associates to each node *v* in a connected weighted graph the positive value *e*(*v*) such that

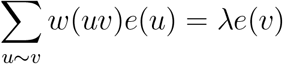

holds for all nodes simultaneously with the largest possible *λ*. In other words, 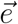 is the Perron-Frobenius eigenvector of the graph’s adjacency matrix. Since this vector is only defined up to a constant, we use the default normalization, which takes it to be a unit vector in the Euclidean norm. We show an alternative normalization, which makes the minimum entry 1, in Figure 1 (d).

Eigenvector centrality is closely related to the adjacency matrix-based spectral methods we discuss below. Since the adjacency matrix of a phylogenetic tree with *n* tips has exactly 4*n* − 4 non-zero entries, it follows that eigenvector centrality can also be computed to a fixed precision *ϵ* in linear time, with a constant proportional to log(1*/ϵ*). It can also be shown - as we do in Section 6.7 in the **Supplementary Materials** - that the largest value of eigenvector centrality in a phylogenetic tree is achieved by an internal node, though it will not necessarily be the root (this result holds for trees with arbitrary positive branch lengths). The distribution of this quantity, maximum eigenvector centrality, for unweighted (unit branch length) phylogenetic trees with *N* = 45 (*n* = 23), is shown in Figure S15 in the **Supplementary Materials**.

#### 3.4.6 Spectral properties of trees

Spectral properties are properties of the eigenvalues of matrices associated with graphs, such as the adjacency, Laplacian or normalized Laplacian matrices [20]. Since spectral methods are a key tool of algebraic graph theory, it is natural to ask whether they can capture important features of pathogen phylogenies. Some of the eigenvalues are interpretable in a limited number of cases [20]. The eigenvalues of the Laplacian of a graph reveal its number of components, the number of spanning trees it has, and the minimal energy of its balanced orthogonal representation in ℝ^*m*^. The second smallest eigenvalue provides a bound on the *conductance* of the graph, the minimum ratio of the number of edges crossing a cut to the size of the cut. The eigenvalues of the normalized Laplacian also provide information on the conductance of a graph as well as its diameter, number of bipartite components and properties of random walks defined on it [9].

Since trees are bipartite graphs, the spectra of their adjacency matrices are symmetric around 0 [20] and the spectra of their normalized Laplacians are symmetric around 1 [9]. The full set of eigenvalues of a tree’s adjacency matrix reveals the number of *matchings* (node-disjoint collections of edges); namely, the characteristic polynomial, which has these eigenvalues as its roots, has coefficients whose absolute values equal the number of matchings of each size [42].

#### 3.4.7 Spectral properties derived from the distance Laplacian matrix

In a recent paper, Lewitus and Morlon [36] used the spectra of the distance Laplacian matrix, obtained by subtracting the distance matrix from a diagonal matrix formed by its row sums, to characterize and compare trees. They found promising results and tentatively stated that these spectra were likely to distinguish trees from one another. In Section 6.8 of our **Supplementary Materials**, on the negative side, we exhibit a pair of trees on *n* = 3 tips with different branch lengths that nevertheless have identical distance Laplacian spectra. On the positive side, we verified that no pair of different phylogenetic trees with at most 32 tips and unit branch lengths has this property; however, we suspect that, as has been shown for most other spectra [42], the fraction of trees on *n* tips uniquely defined by their distance Laplacian spectra tends to 0 for large *n*.

We use the four summary statistics proposed by the authors [36] - namely, the maximum eigenvalue, and the asymmetry, kurtosis, and maximum density of the eigenvalue distribution (obtained via smoothing with a Gaussian kernel), as implemented in the *RPANDA* [48] package in R [59]. We note that, unlike the rest of the statistics in this paper, these ones require a number of operations proportional to *n*^3^, where *n* is the number of tips, and thus take substantially longer to compute.

## 4 Results

### 4.1 Tree topology differentiates viruses and epidemiological scenarios

We find that tree topology carries considerable information and differs both between viruses and between different epidemiological scenarios for the same virus, on real as well as simulated data. While degree sequence, clustering coefficients and other measures based in network science are not informative for binary trees, a number of non-standard topological features differ. Furthermore, there are several features that distinguish well between groups of trees with the same overall level of asymmetry, highlighting the need to move beyond asymmetry when using trees to infer evolutionary and epidemiological parameters or to test hypotheses [23]. Figures 2 and 3 illustrate the distributions of all tree statistics considered in this paper. In addition, we perform a (two-sided) Mann-Whitney U test, also known as a Wilcoxon rank-sum test [39], between each pair of scenarios or viruses in a given group, for every statistic we consider, and report the *p*-values in Tables S2 and S3 in the **Supplementary Materials**. We then apply a Bonferroni correction [15] to account for the multiple hypotheses being tested, and highlight in bold those values which remain significant at the *α* = 0.05 level after this correction.

**Figure 2:**
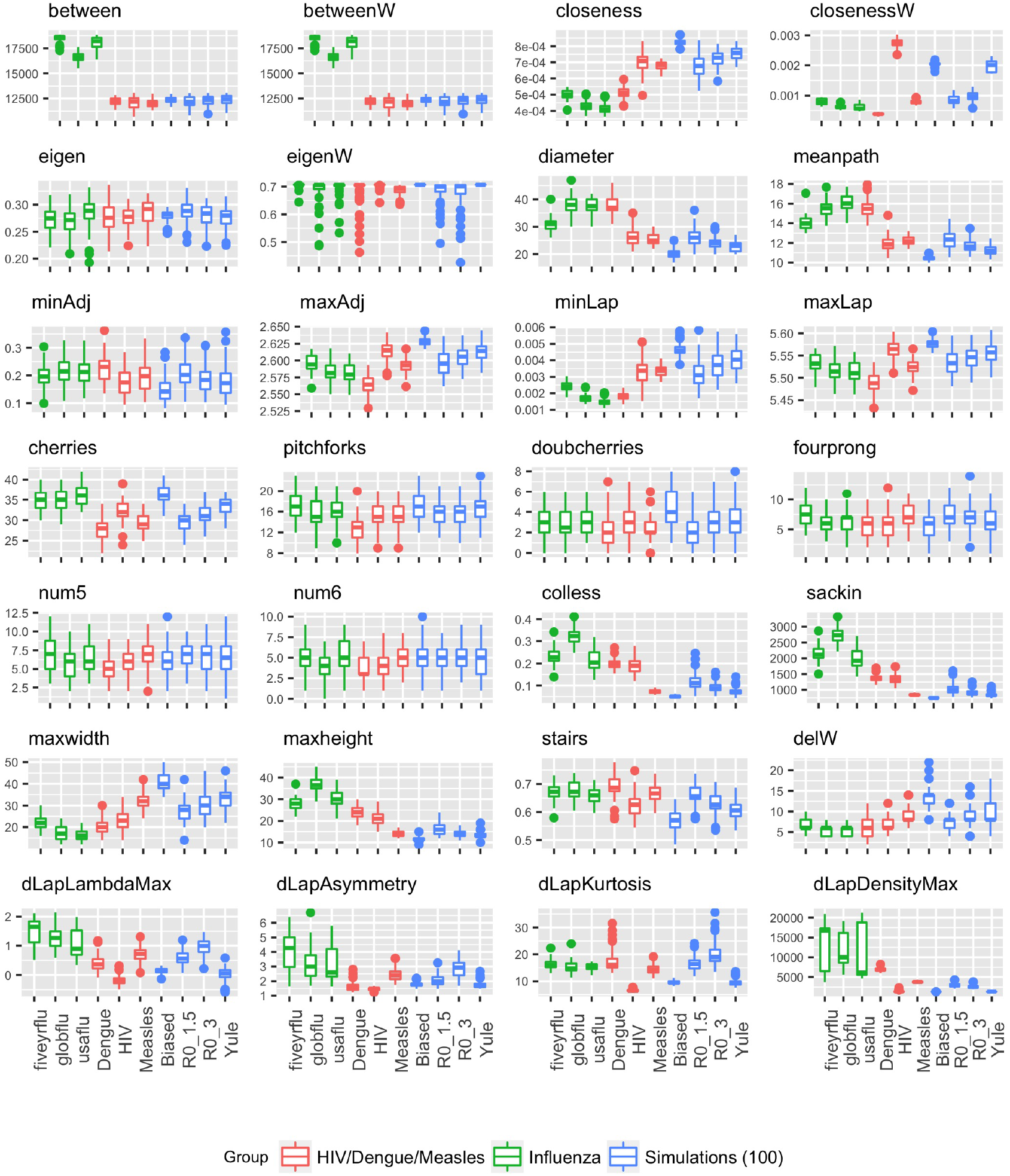
Tree summaries for the HIV/Dengue/Measles, influenza and simulations of size 100

**Figure 3:**
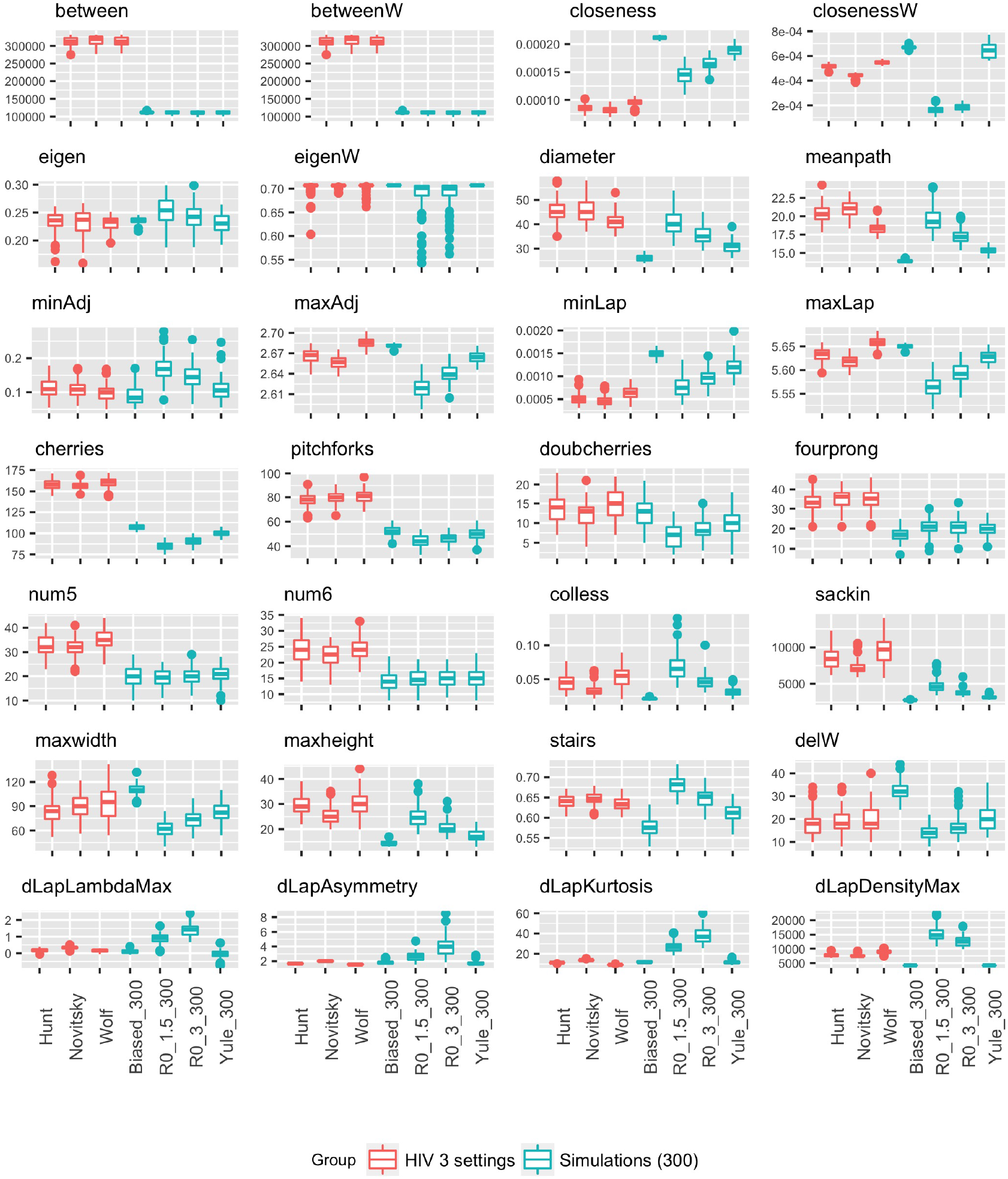
Tree summaries for HIV in three settings and simulations of size 300

Most of the statistics do not vary much between the three viruses in Figure 2 (red boxplots). Distinguishing the topologies in these groups of trees requires tools going beyond the traditional symmetry or imbalance metrics; in this case, the only statistically significant difference are produced by the number of cherries, weighted closeness centrality, maximum height, and the proportion of imbalanced subtrees (stairs) capture differences that are not apparent in the imbalance. In contrast, while two of the spectral statistics (maximum adjacency and maximum Laplacian eigenvalues) show statistically significant differences among the groups, the vertical scale is very small, and these differences are a small percentage of the overall value (2% for the maximum adjacency and 1% for the maximum Laplacian).Furthermore, three of the network statistics - namely, the Wiener index (meanpath) and the weighted closeness and eigenvector centralities - also distinguish the three viruses, as do three of the four statistics based on the distance Laplacian.

The green boxplots in Figure 2 show the tree summaries for influenza in three scenarios. It is well-known that global influenza patterns give rise to highly imbalanced trees, an observation which in part motivated the growing field of phylodynamics [23]. In our data, the global five-year and (2-year) USA flu trees have similar, lower, levels of asymmetry. The only statistics which are able to differentiate all three groups at a statistically significant level after multiple testing correction are betweenness centrality and the maximum Laplacian eigenvalue. As with the three viruses, the differences in the latter are not pronounced, and are very small in relative magnitude. In addition, the LM spectral statistics are quite discriminating for these scenarios, though none of them is able to do so in a statistically significant way. Figure 6 shows a random tree from each group, and differences between the shapes of the flu trees are not immediately apparent by eye.

The blue boxplots in Figure 2 show the summaries for small trees (100 tips) in the four simulated settings. This time, only the closeness centrality, the diameter, and the mean path, as well as the number of cherries, the Sackin and Colless imbalances, the maximum height, the stairs statistic and the lambdaMax statistic based on the distance Laplacian spectrum, are able to differentiate every pair of these scenarios in a statistically significant way.

In contrast, the cyan boxplots in Figure 3 show that large trees in the same simulated settings can be discriminated by all of these statistics, but also by several other ones, including the adjacency and Laplacian eigenvalues as well as the maximum width and the maximum difference in width (delW). This suggests that it is strictly easier to discriminate larger trees than it is to discriminate smaller trees.

Lastly, the three epidemiological scenarios (concentrated epidemic, generalized epidemic in a village, and generalized epidemic in a country) for HIV, shown in the red boxplots in Figure 3, appear to be quite difficult to distinguish. Only the weighted and unweighted closeness, as well as the Sackin and Colless imbalance, the maximum eigenvalues of the adjacency and the Laplacian, and the asymmetry and kurtosis of the distance Laplacian spectrum, are able to do so in a statistically significant manner.

Figure 4 shows the mutual information between the virus or epidemiological scenario and the value of each statistic, for the 5 settings we tested. We have computed these values by discretizing the range of each tree shape statistic into 20 equally-sized bins. The first panel shows that the highest mutual information with the virus (HIV, Dengue or Measles) is obtained by a network science-based statistic in Figure 4, namely, the weighted closeness, with the densityMax coming close second, while the second panel shows that the highest mutual information with the type of flu (global, US or 5-year) is obtained by the minimum Laplacian eigenvalue, with the betweenness (which is the same whether it is weighted or unweighted) and densityMax coming close behind. In the remaining three scenarios (third, fourth and fifth panels), closeness centrality is the only statistic consistently in the top 3 highest mutual information with the model category or scenario.

**Figure 4:**
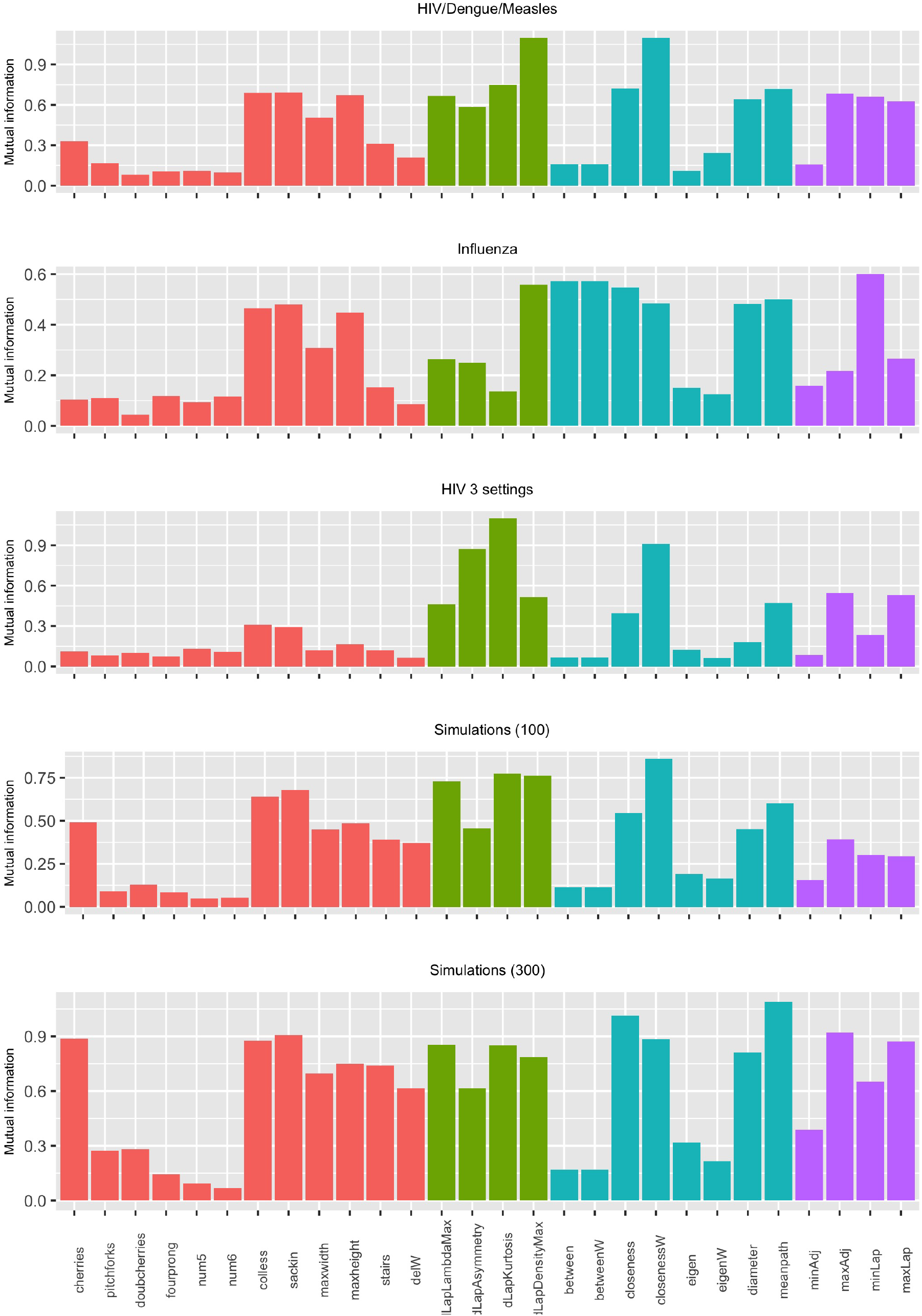
Mutual information between tree summaries and the virus type or epidemiological scenario. Labels are as in Table 1, and the colour indicates the type of statistic as per Table 1, i.e. red: basic statistics; green: distance Laplacian spectrum statistics; blue: network science-based statistics; purple: spectral statistics.

### 4.2 The addition of network statistics improves classification

In some scenarios, the trees are markedly different and only a few features are necessary to distinguish them. For example, in the influenza scenario it is clear from Figure 3 that the Colless, Sackin, betweenness and maximum heights statistics all distinguish between global influenza and the other two groups, whereas the closeness measures, the diameter, the mean path and the maximum width distinguish the five year trees from the other two groups. Asymmetry measures alone do not distinguish the three groups.

We use a generalised linear model and two flavours of random forests to classify trees within each scenario. For example, we classified trees as HIV, Dengue or Measles; we classified influenza trees as five-year, global, and USA; we classified simulated trees as biased, Yule, *R*_0_ = 1.5 or 3; and we classified the HIV trees by epidemiological scenario. In each classification task, we randomly selected 75% of the trees (75 trees from each group) for training and used the remaining 25 trees from each group for testing, and computed the classification error of the predictor on the testing set. We report the overall classification error with and without the features based on network science, as well as with and without the features based on the distance Laplacian spectrum (abbreviated as “LM” statistics after the authors Lewitus and Morlon). The results are shown in Figure 5 (a). It is clear that the standard (“basic”) tree shape statistics are not able to get close to perfect classification on any of the datasets except for the (relatively simple) task of distinguishing three different viruses. The addition of the costly LM statistics improves the performance, but so does the addition of the easily computable new network science-based statistics we introduce in this paper, with comparable gains in performance relative to the baseline (a larger improvement on the flu scenarios, and a slightly smaller one on the HIV scenarios and on simulated data). Interestingly, the addition of network statistics actually increases the error relative to the baseline on the large simulated trees, an effect that is likely due to the random nature of the classifier.

**Figure 5:**
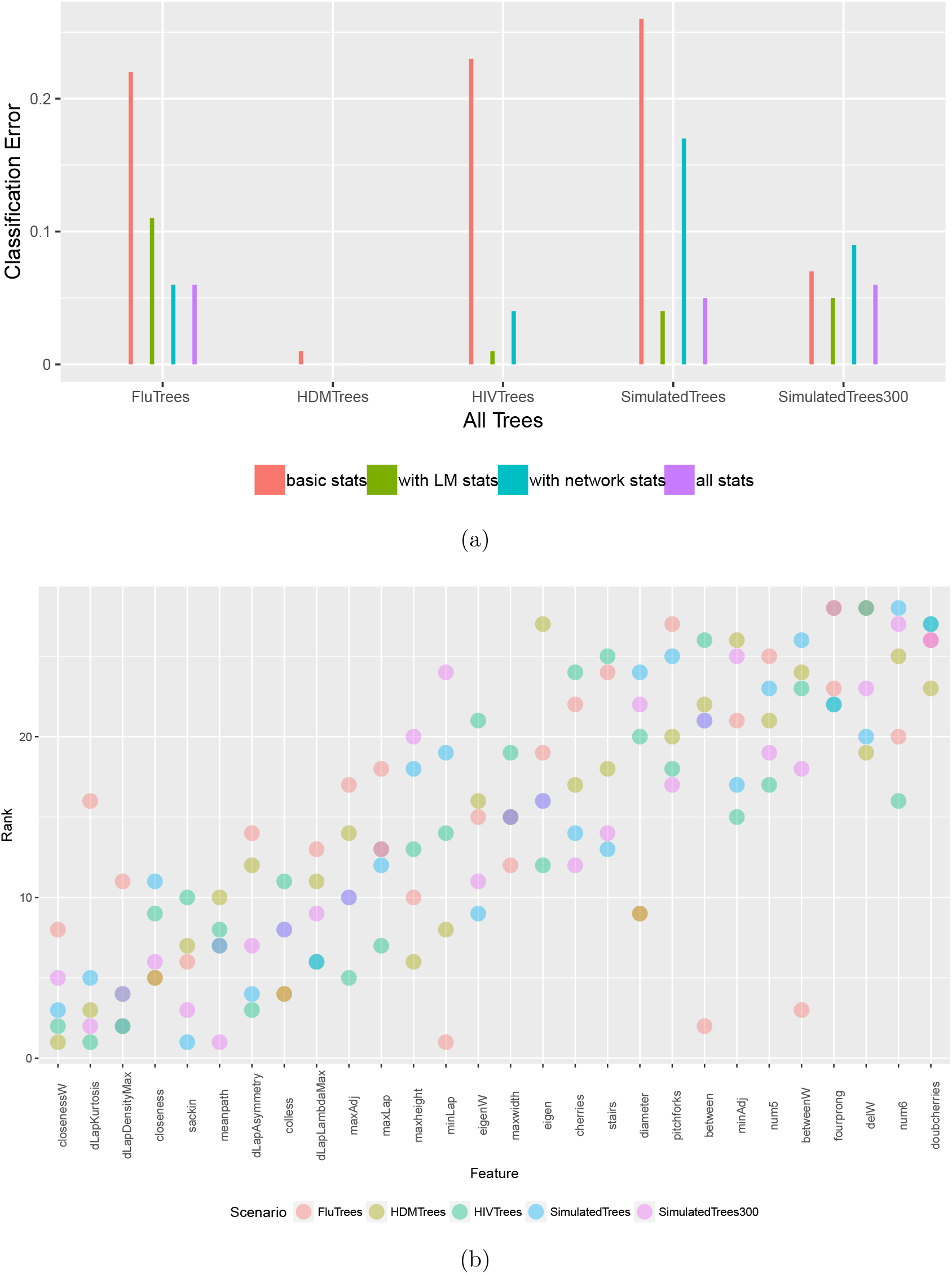
(a) Accuracy of classification with and without network science features, as well as with and without the distance Laplacian spectral features. (b) Feature importance in multi-class classification across all scenarios, ordered by the median. Here the importance for each feature is computed based on the mean decrease in node impurity (we use *varImp* function [33] in *R* which uses Gini index as an an impurity function). Each point is the rank of the corresponding feature in one of the classification tasks. Low ranks correspond to the most important features (i.e. the top-ranked feature has rank 1, and so on).

**Figure 6:**
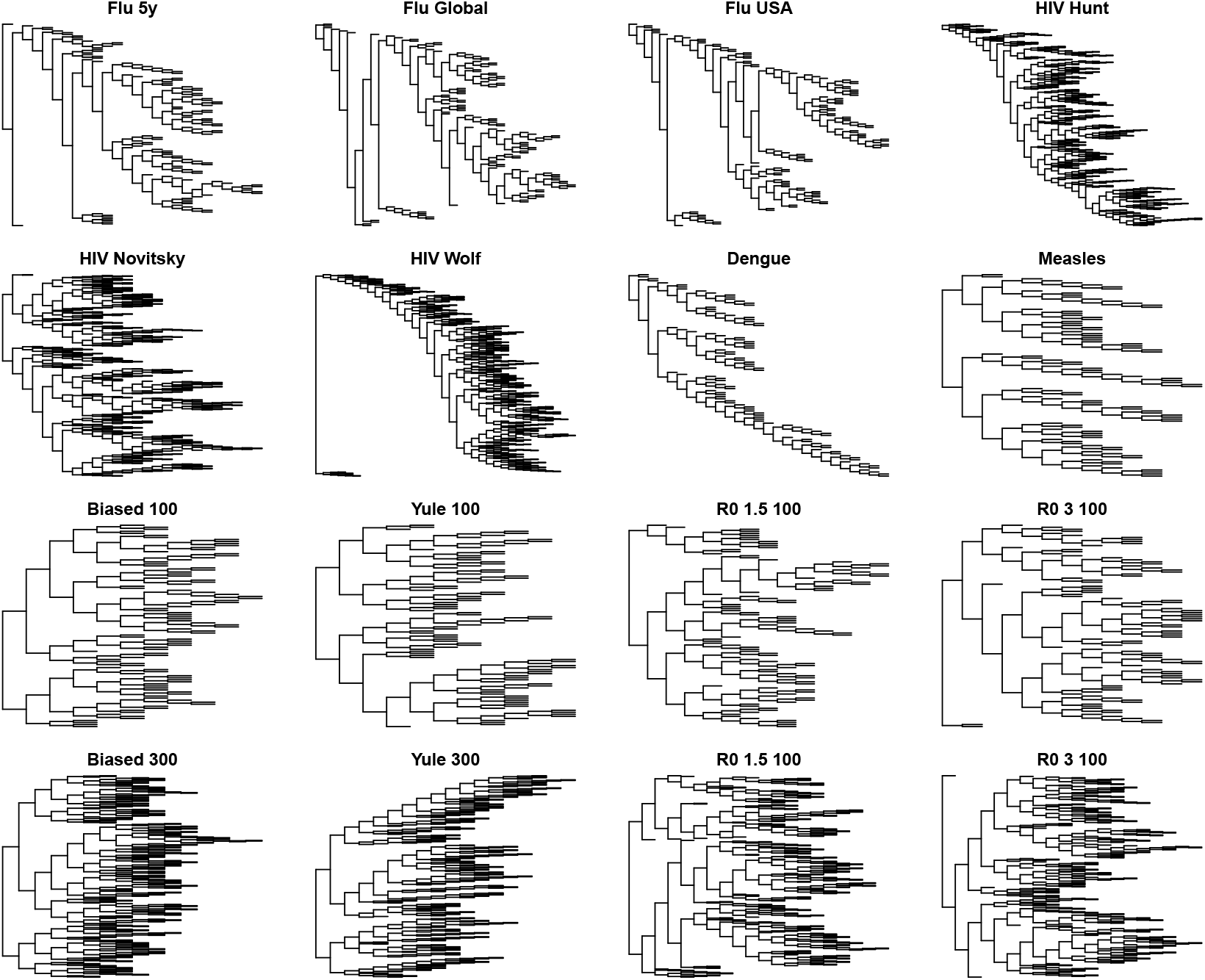
A randomly sampled tree from each scenario (except HIV in the 3-virus comparison because HIV is represented in three other trees). To allow for focus on tree shape rather than on branch lengths, trees have been visualized with branch lengths set to 1.

Both random forests and the GLM regression additionally provide estimates of the importance of each feature. For the random forest classifier, the prediction accuracy on the out-of-bag portion of the data is recorded for each tree, and then the same is done after randomly permuting each predictor variable. The differences between the two accuracies are then averaged over all trees, and normalized by the standard error. For the GLM regression, the absolute value of the *t*-statistic for each model parameter is used as the importance measure. We used the *caret* package [34] in R [59] to compute these for all the classifiers. For each classification task in each scenario, we then ranked the features by importance. We show the rank data in Figure 5 (b). It is apparent that weighted closeness centrality is the feature with the highest importance (lowest rank) across all scenarios, on average, with the much costlier to compute distance Laplacian-based features coming close behind.

## 5 Conclusions

We have used network science to developed tailored statistics to capture the topologies of phylogenetic trees and illustrated how they compare across different viruses and scenarios. The viruses vary in pathogenesis and their rates and modes of transmission, characteristics which ideally should be reflected in the tree shape. We find that network-science-derived statistics are complementary to asymmetry and frequency of small configurations with a tree. We have also used a tree kernel to compare trees; while it distinguishes well between three different viruses, the statistics do a better job of distinguishing the three influenza scenarios. However, no individual statistic is sufficient to consistently capture the diversity of tree topologies.

It is highly likely that investigators will need to use multiple summary measures to resolve biologically meaningful differences among phylogenies. This is a problem of *feature selection*. Indeed, some of the literature on phylogenetic tree topologies has focused on evaluating which summary measures are the best suited for detecting informative differences in tree topologies [31, 47, 1]. This effort makes the implicit assumption that there exists a specific subset of features that will always be a good choice for all data sets (or data for a given domain), which is not necessarily the case. There may exist summary measures that have not been formulated but would confer an optimal fit to a given dataset. Matsen [41] proposed an innovative approach to this problem, where a generalized and flexible class of tree topology statistics (which can express many established measures of tree shape) can be adapted to the data using a genetic algorithm.

The underlying evolutionary or epidemiological process may affect tree topologies, but they are likely also to be affected by sampling density and uniformity, geographical constraints and population structure, ecological dynamics and host contact structure (in the case of pathogens). Our results point to the important role of tree topologies as a vehicle for obtaining information about some of these underlying processes. Topological summary features are readily interpretable, easy to compute and scale readily to large trees, even when they do not necessarily provide direct insights into the underlying evolutionary or diversification processes. The network-based and spectral statistics we have described also generalize naturally to phylogenetic networks.

On the other hand, each feature can capture only a narrowly-defined aspect of tree shape, such as imbalance. This could potentially be overcome by using multiple features, although correlations among features may lead to diminishing returns with the addition of new features. Features can also be difficult to compare among trees of different sizes (numbers of tips), and despite attempts to derive normalized statistics they remain sensitive to size. However, Pompei and colleagues [55] have proposed a randomization procedure to derive the null distribution of summary features as a possible resolution of this problem.

## Acknowledgments

This work was in part supported by the AWS Cloud Credits for Research program. The authors would like to thank Nithum Thain and Stepan Grinek for providing computational support to this project, and Kamyar Khodamoradi for pointing out the theorem by Jordan relevant to the existence of centroids. LC was supported by an NSERC Discovery Grant (RGPIN-2016-04622) as well as an Alfred P. Sloan Fellowship (FG-2016-6392). LC also acknowledges funding from the MRC Centre for Global Infectious Disease Analysis (reference MR/R015600/1), jointly funded by the UK Medical Research Council (MRC) and the UK Foreign, Commonwealth & Development Office (FCDO), under the MRC/FCDO Concordat agreement and is also part of the EDCTP2 programme supported by the European Union. CC was supported by the Engineering and Physical Sciences of the United Kingdom (EP/K026003/1 and EP/N014529/1). AP was supported by an NSERC Discovery Grant (RGPIN-2018-05516) as well as a CIHR Project Grant (PJT-155990). This research was undertaken, in part, thanks to funding from the Canada 150 Research Chairs Program (CC).

## 6 Supplementary Materials

### 6.1 GenBank accession numbers for outgroups

**Table S1:** Accession numbers for outgroups

| Virus | Outgroup | GenBank accession number |
| --- | --- | --- |
| HIV-1 subtype B | a subtype D sequence | AY071949 |
| Dengue virus serotype 4 | isolate from the Philippines (1956) | U18433 |
| Measles virus | a genotype D6 sequence | AY523581 |

### 6.2 *p*-values for the statistics used to compare viruses or scenarios

The *p*-values come from the Mann-Whitney U test for each statistic in a pairwise comparison; the values passing Bonferroni correction for multiple testing at the *α* = 0.05 level are in bold.

**Table S2:**
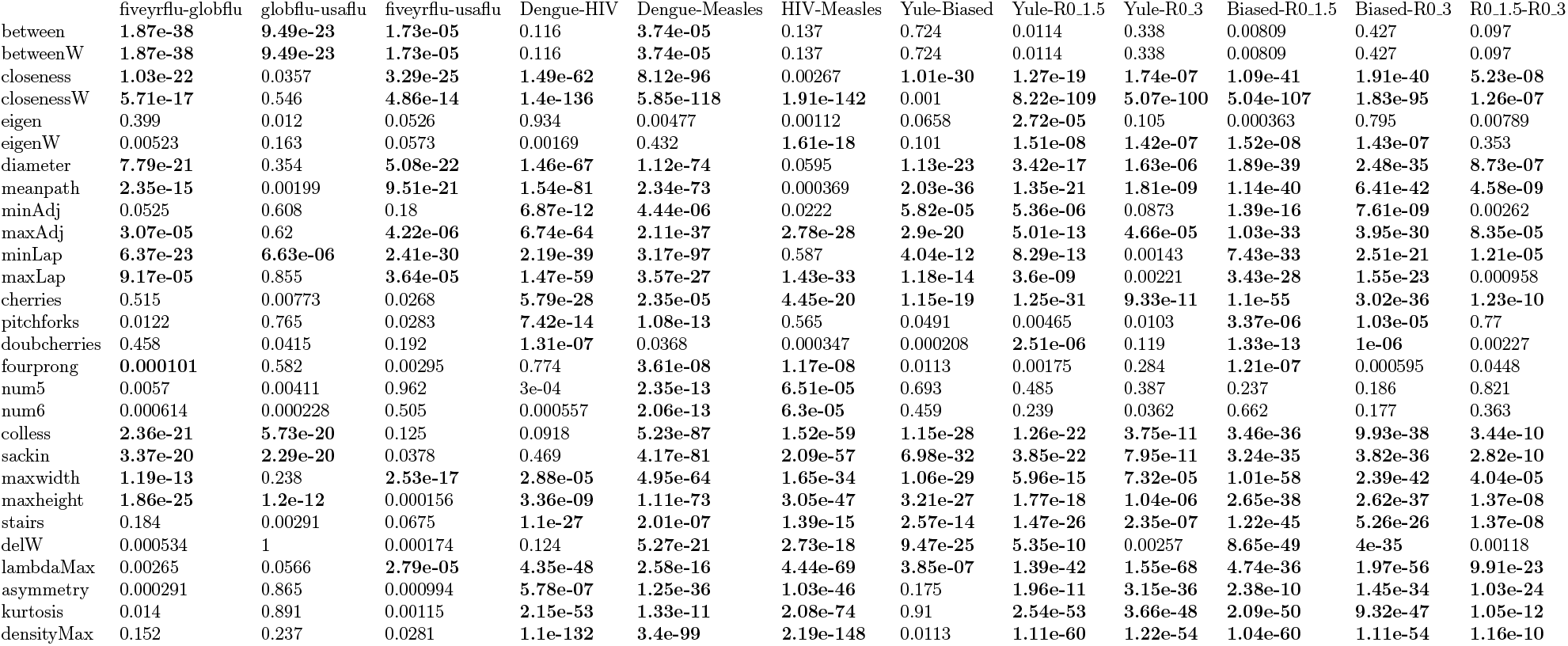
*p*-values for pairs of scenarios in Figure 2.

**Table S3:**
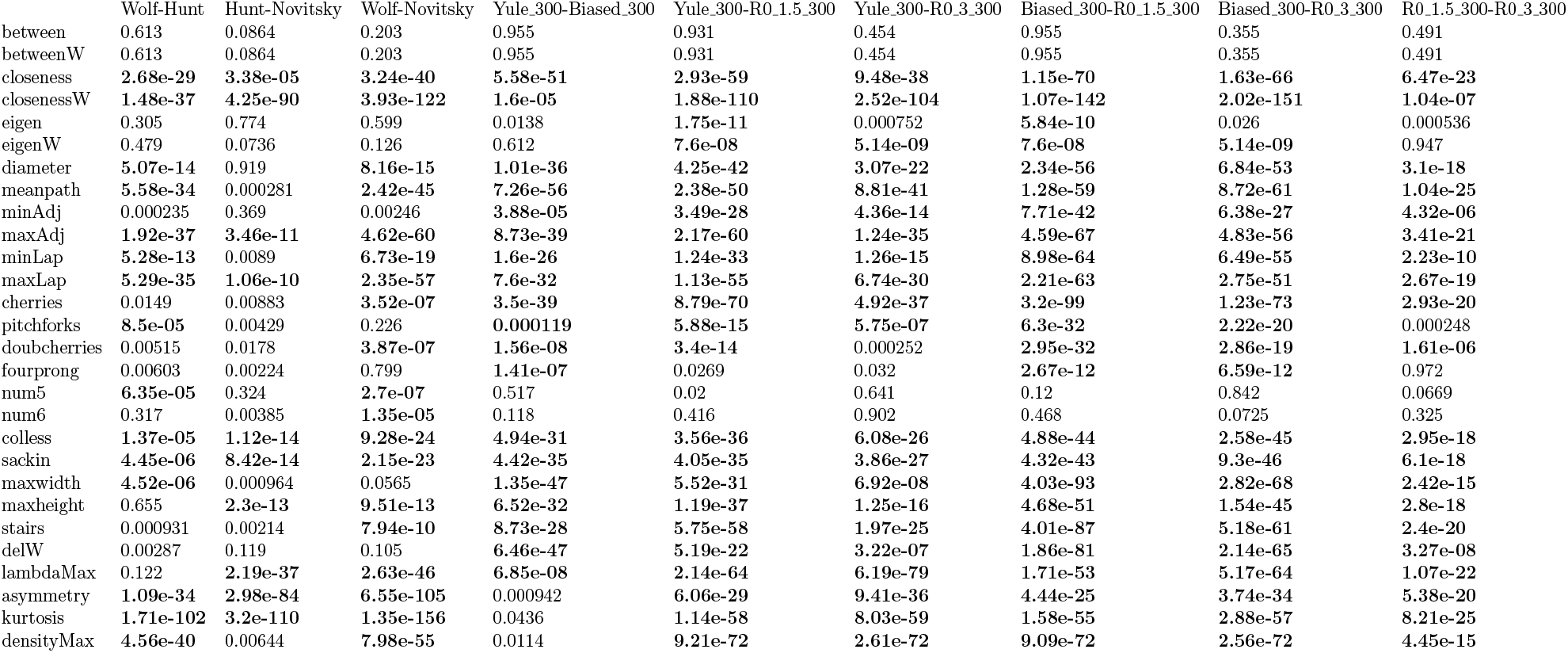
*p*-values for pairs of scenarios in Figure 3.

### 6.3 Kernel-based tree comparisons

Rather than using summary statistics to quantify differences in tree topologies, we can calculate a similarity or distance measure between trees. For example, the Robinson-Foulds distance measure is an edit distance between trees with the same labels (*i.e.*, relating the same taxa). The resulting distance matrix for a set of tree topologies can then be interpreted as a measure space for supervised or unsupervised classifiers [26]. However, the requirement for shared labels in the Robinson-Foulds distance prevents its application to the present problem of comparing phylogenies from different viruses.

To overcome this limitation, we previously adapted a kernel method from natural language processing [13] to provide a similarity measure that operates on both the topology and branch lengths of trees [57]. Every tree is comprised of a large number of subset trees that can each act as a feature. A subset tree is a contiguous set of branches rooted at an internal node of its parent tree, which does not necessarily include all descendants of that internal node. In other words, the subset tree does not have to extend out to the tips of the parent tree.

A subset tree is completely defined by its branching order if we rotate branches of the tree (so-called “ladderization”) so that all branching events occur preferentially to one side. For a tree of even modest size, the number of all possible subset trees is extremely large; the space of all possible subset trees is even more immense. Clearly, it is not feasible to exhaustively enumerate the appearance of every possible subset tree for a given observed tree.

The kernel trick is a well-established technique in machine learning that provides an efficient way to compare trees with respect to their subset trees by limiting the comparison to those features that appear in one or both trees [57]. Calculating the inner products over this restricted set of features for every pair of trees yields a distance matrix which defines a projection of the trees into a high dimensional space with convenient properties for machine learning.

We applied this kernel method to the data sets examined above with the following parameter settings: decay factor *λ* = 0.2; radial basis function variance *σ* = 2.

Figure S1 represents a set of principal component analysis (PCA) plots that illustrate the separation of HIV, Dengue, and Measles virus phylogenies into distinct clusters in the space defined by the tree kernel. The HIV trees are separated from the others along the first principal component that explained roughly 75% of the overall variation. Dengue and Measles virus trees can only be separated into distinct clusters by the third principal component. For comparison, the first two principal components of the tree shape statistics shown in Figure S2 are already able to separate the three viruses. The dots represent individual trees; the arrows point in the direction of maximum multiple correlation with the principal components for each tree shape statistic, while their length indicates the strength of this multiple correlation. Once again, we see that the network science-based statistics are overrepresented among the longest arrows.

Furthermore, applying the same method to trees from the three different samples of influenza A virus outbreaks was less promising (Figure S4). While it is possible that adjusting the kernel tuning parameters (*λ* and *σ*) could yield better results, we would risk over-fitting this method to these data. In contrast, Colless and Sackin imbalance separate global flu from the other two groups, while closeness, diameter and mean path separate the five-year flu trees from the global and USA groups. A combination of these summary statistics separate the three groups, as shown in Figure S3 below.

We observed a similar situation in the case of the other three datasets, in which the first three principal components of the kernel-based method are sometimes needed to separating the different categories or scenarios (Figures S5, S7 and S8, while only the first two principal components of the tree shape statistics considered in this paper are usually sufficient (Figures S6 and S9). However, both methods are clearly able to identify enough signal in the data to separate the categories or scenarios.

**Figure S1:**
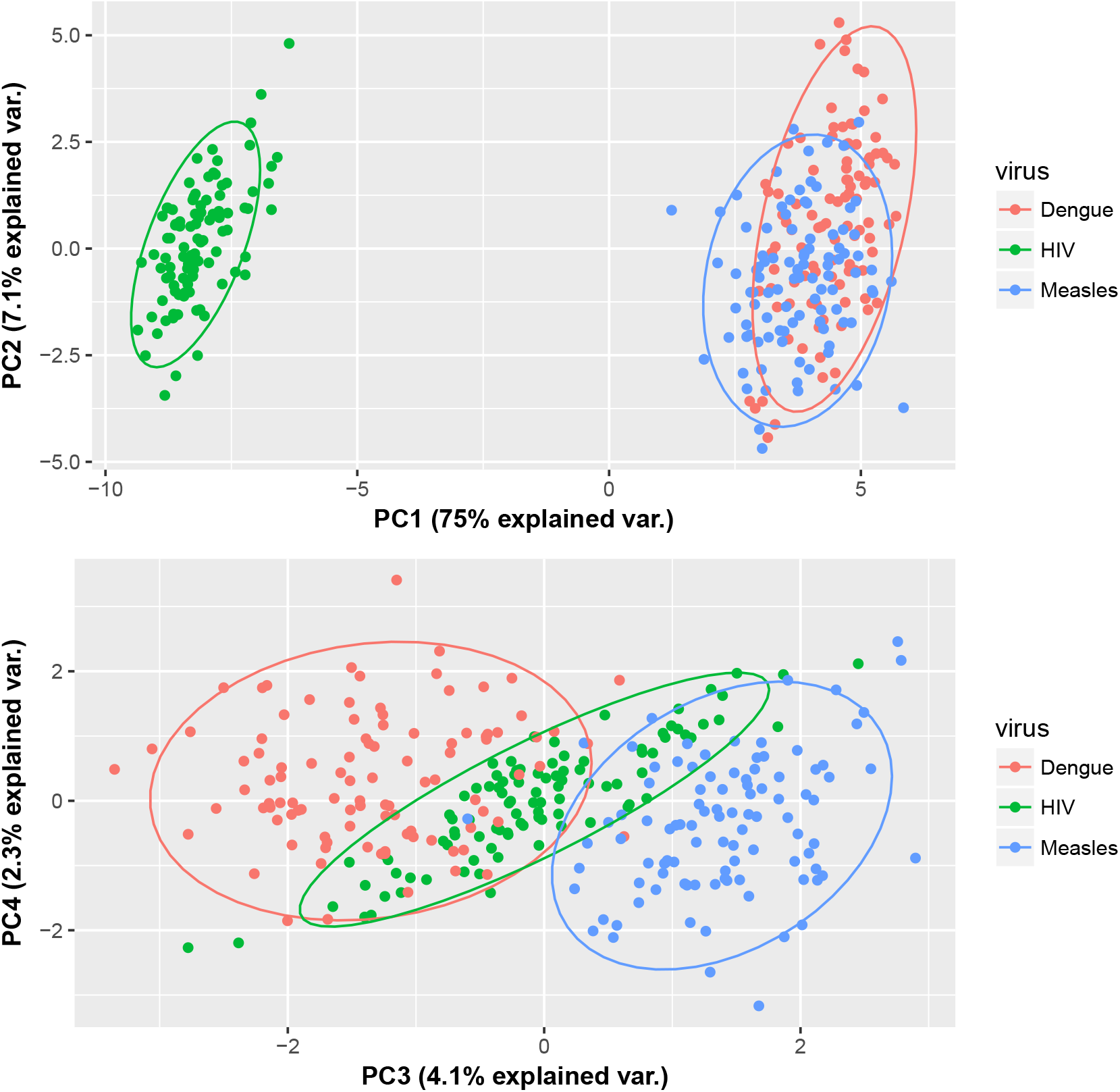
Kernel PCA plots of first four principal components illustrating the separation of HIV (green), Dengue (red) and Measles (blue) virus phylogenies by a tree kernel method.

### 6.4 Calculating the (degree) assortativity of a phylogenetic tree

Consider a phylogenetic tree on *n* tips as a graph. Its degree assortativity is the Pearson correlation between the degrees of the sources (heads) and targets (tails) of its edges. We direct each edge away from the root, towards the tips. We show below that for *n* ≥ 4, only two values of the assortativity coefficient can occur (only one such value can obviously occur for *n* ≤ 3).

We make use of the following formula for the Pearson correlation between 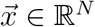 and 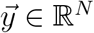:

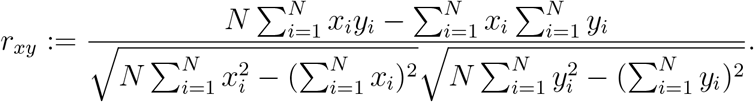

Based on this formula, we note that only two options can occur; the first option is when the root’s children are both internal nodes, and the second option is when one of its children is a tip and the other, an internal node (which occurs when, for instance, there is an outgroup in the data). In the first case, we use

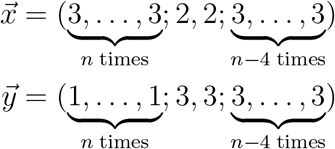

since there are exactly *n* edges going from an internal node to a tip, 2 edges from the root to an internal node, and *n* − 4 edges from an internal node to another internal node. An easy calculation then yields

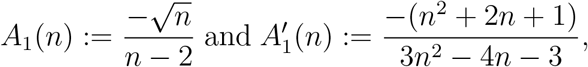

where *A*_1_ denotes the *directed* degree assortativity, calculated by substituting 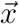 and 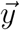 directly into the formula above, while 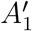 denotes the *undirected* degree assortativity, calculated by substituting 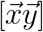 and 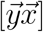 into it, where 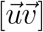 denotes the concatenation of two vectors. The latter occurs because in the undirected case, each directed edge (*ij*) is counted twice, once as an edge from *i* to *j* and once as an edge from *j* to *i*.

**Figure S2:**
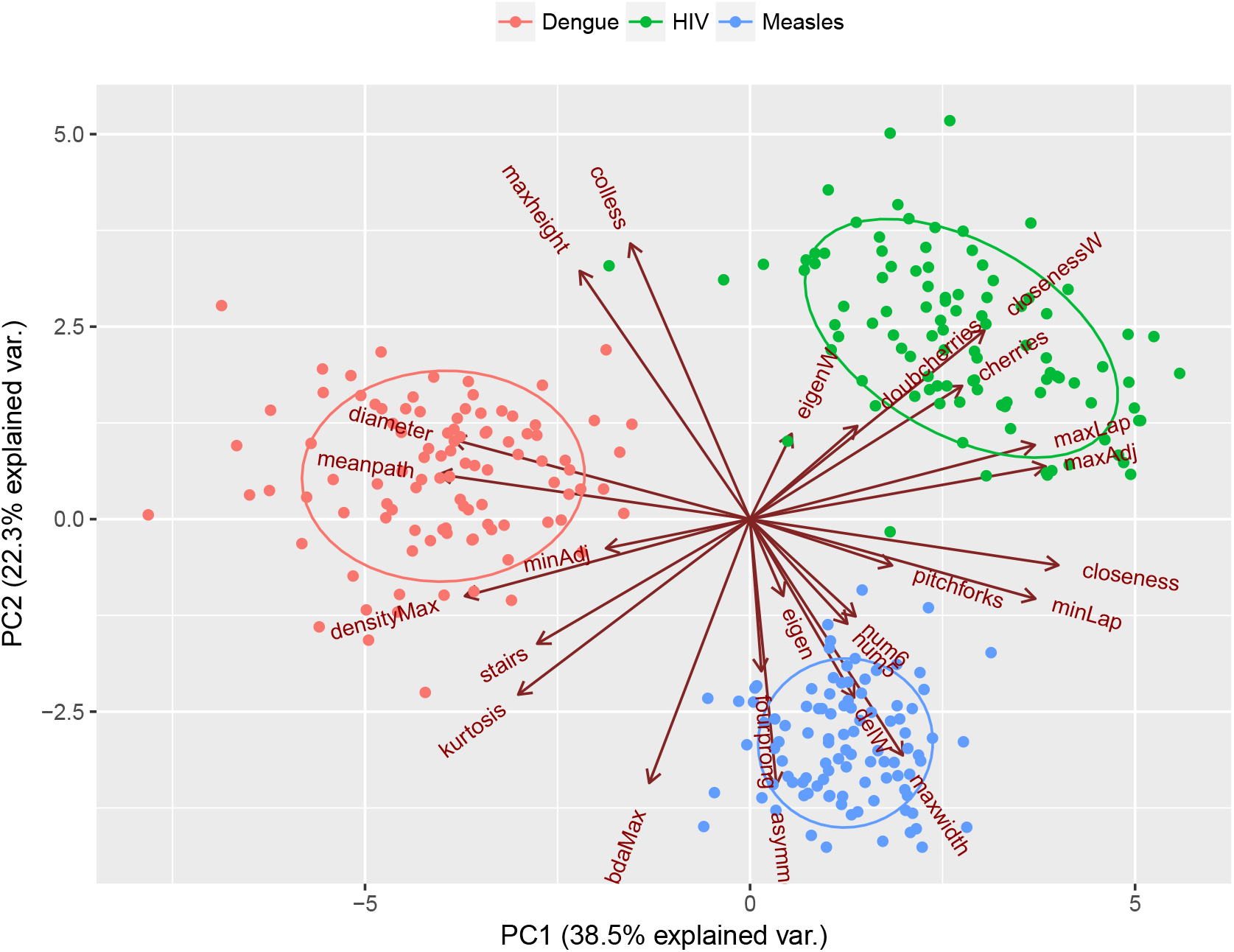
PCA biplots of first two principal components illustrating the separation of HIV (green), Dengue (red) and Measles (blue) virus phylogenies by the shape statistics.

In the second case, where the root has a tip as one of its children, the degree sequences are

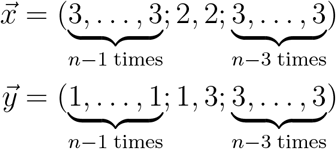

and the corresponding values are

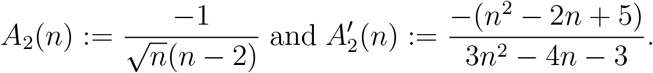

We note that lim_*n*→∞_ *A*_1_(*n*) = 0 = lim_*n*→∞_ *A*_2_(*n*) and 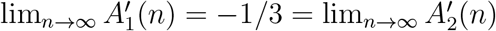.

**Figure S3:**
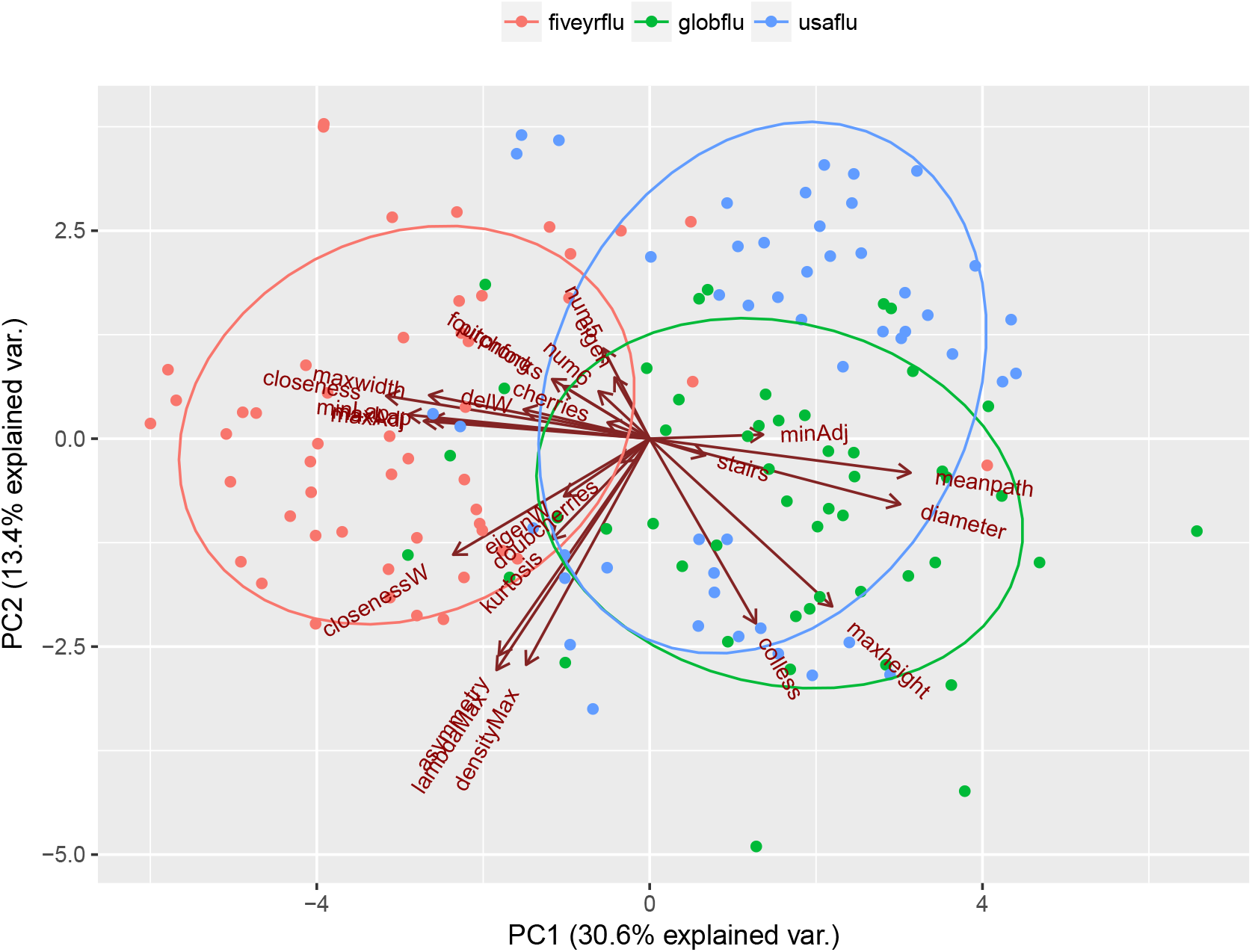
PCA biplots illustrating the separation among influenza A virus phylogenies sampled at global (green), five-year (red) and regional (United States, blue) scopes by using tree shape statistics

### 6.5 The distribution of the diameter and the Wiener index of trees

The distribution of diameters for *n* = 23 tips ranges from 8 to 23, with a mean of 14.147. Figure S10 shows the distribution of all the diameters, together with one of the trees achieving the minimum and maximum values of the diameter, respectively.

The distribution of Wiener indices for *n* = 23 tips ranges from 5382 to 8778, with a mean of 6516.541. Figure S11 shows the distribution of all the diameters, together with the trees achieving the minimum and maximum values of the Wiener index, respectively.

Figure S12 illustrates all 6 phylogenetic trees on *n* = 6 tips with their Wiener index and diameter, arranged in increasing order of the Wiener index, with ties broken by the diameter.

### 6.6 The distribution of betweenness/closeness/eigenvector centrality

The distribution of maximum betweenness centrality values for *n* = 23 tips ranges from 505 to 645, with a mean of 587.592. Figure S13 shows the distribution of all the maximum betweenness centrality values, together with one of the trees achieving the minimum and maximum values, respectively.

The distribution of maximum closeness centrality values for *n* = 23 tips ranges from 143 to 285, with a mean of 189.920. Figure S14 shows the distribution of all the maximum closeness centrality values, together with the trees achieving the minimum and maximum values, respectively.

The distribution of maximum eigenvector centrality values for *n* = 23 tips ranges from 0.2324 to 0.4336, with a mean of 0.3573. Figure S15 shows the distribution of all the maximum eigen-vector centrality values, together with the trees achieving the minimum and maximum values, respectively.

**Figure S4:**
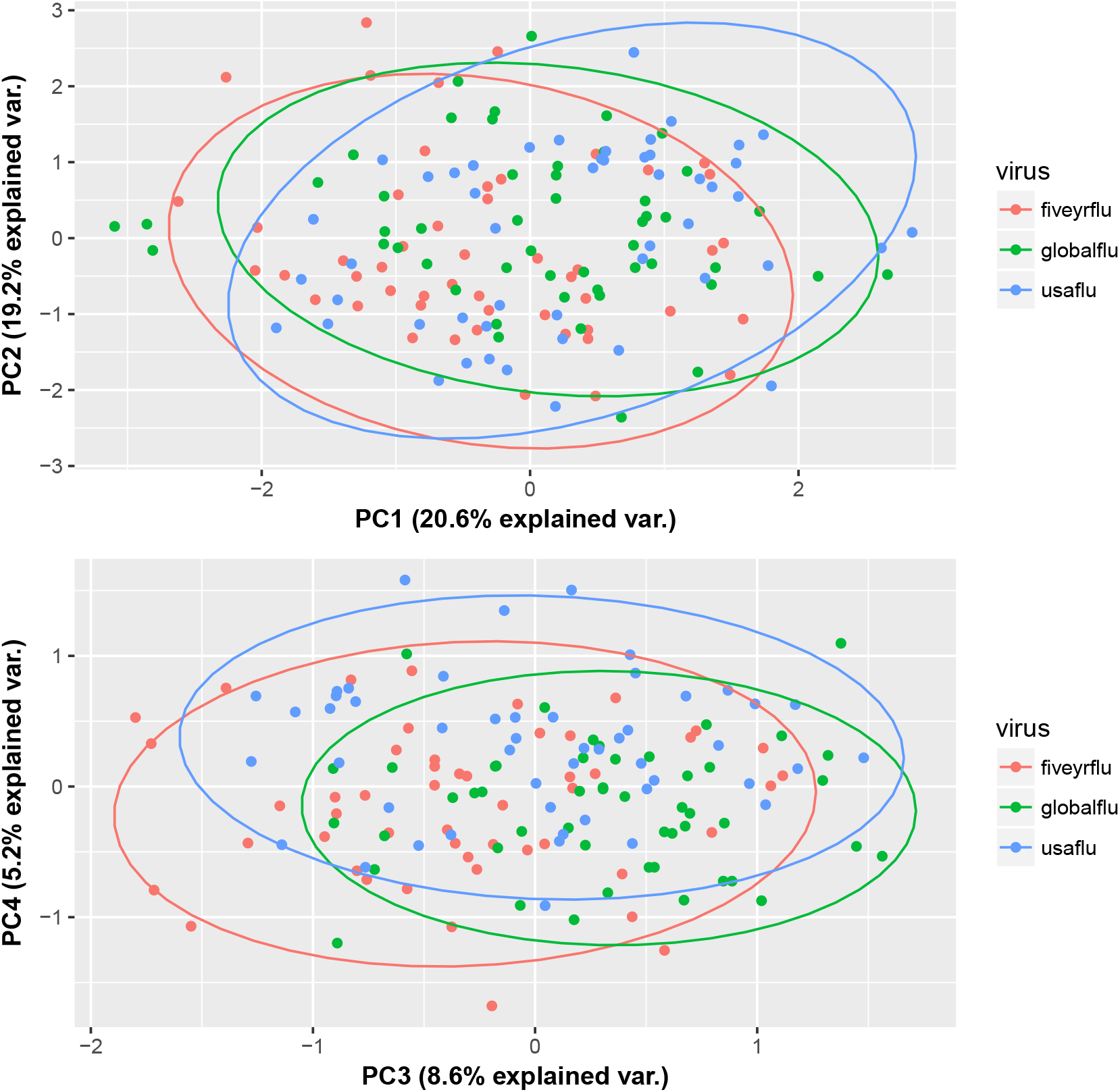
Kernel PCA plots illustrating the overall lack of separation among influenza A virus phylogenies sampled at global (green), five-year (red) and regional (United States, blue) scopes by using the tree kernel method.

Figure S16 illustrates all 6 phylogenetic trees on *n* = 6 tips with their betweenness, closeness, and eigenvector centrality, arranged in increasing order of the betweenness, with ties broken by the closeness.

### 6.7 Proof of the location of the maximum values of centralities

We now present a simple proof for the following facts, mentioned in the main text:

a. Betweenness centrality is maximized at the **tricenter** of the tree (the node which, if the tree is rerooted at it, has 3 descendant clades of sizes within one of each other), if one exists, and minimum at the tips
b. Closeness centrality is maximized at the **centroid** of the tree (the node which, if the tree is rerooted at it, has all descendant clades containing fewer than half the total nodes), and minimum at one of the tips
c. Eigenvector centrality is maximized at one of the internal nodes

**Figure S5:**
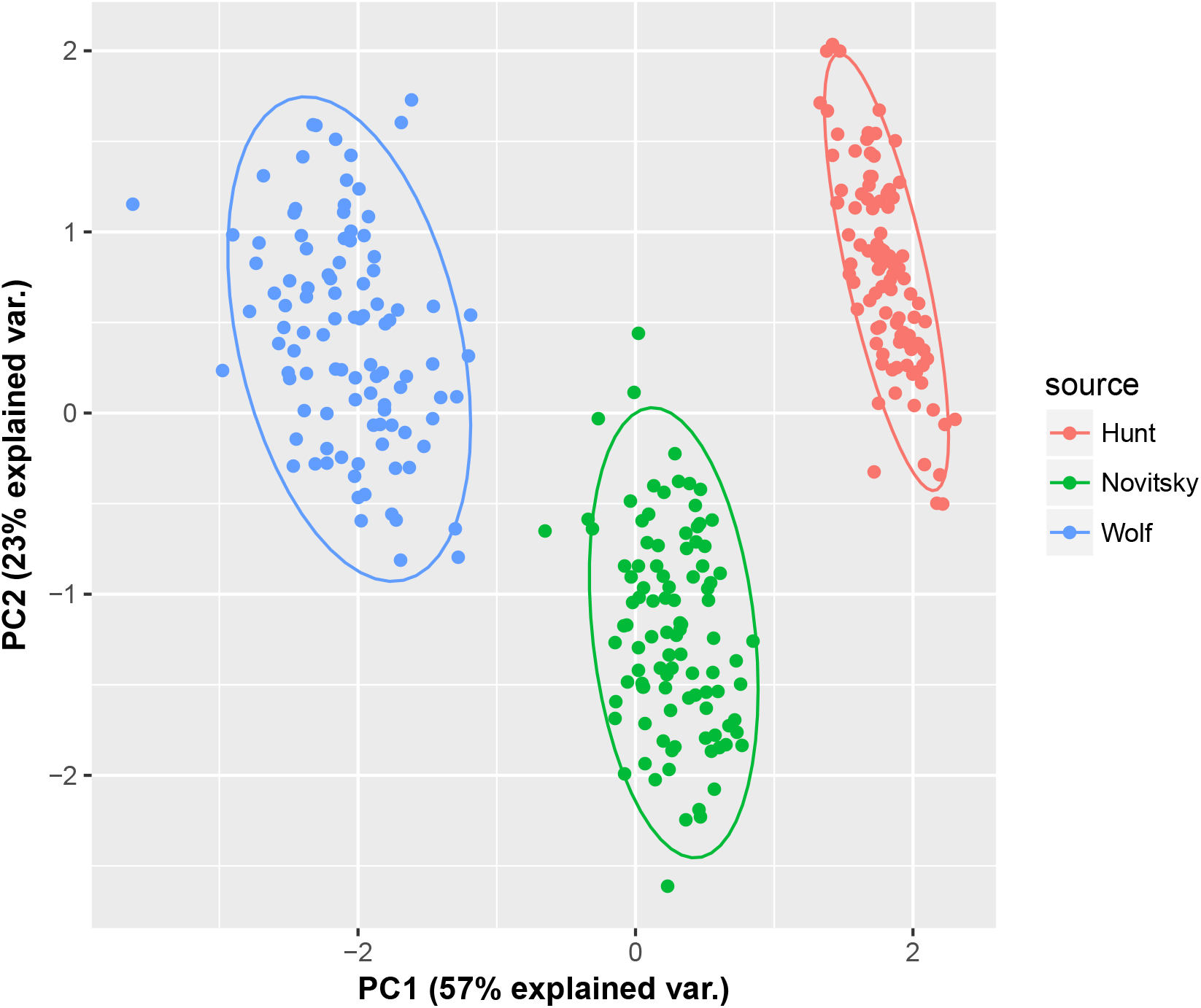
Kernel PCA plots illustrating the separation among HIV phylogenies sampled from a concentrated epidemic (blue), a generalized epidemic in a country (red) and in a village (green) by using the kernel method.

**Proof**:

a. Let *v* be any node in a tree *T*. If *n*_1_ and *n*_2_ are the sizes of the left and right subtrees of *v* and *n*_0_ is the number of nodes outside *T_v_*, the clade of *T* subtended by *v*, then its betweenness centrality is *n*_0_*n*_1_ + *n*_0_*n*_2_ + *n*_1_*n*_2_. We maximize this quantity subject to the constraint *n*_0_ + *n*_1_ + *n*_2_ = *N* − 1 and keep in mind that all of the sizes are integers. First, if *N* ≡ 1 mod 3, all three of these quantities can be equal to 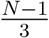, and this matches the best we can do even if we allow fractional values. Second, if *N* ≡ 2 mod 3, say *N* = 3*K* + 2, the best option is to set one of the three values to 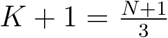 and make the remaining two equal to 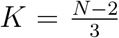. Finally, if *N* ≡ 0 mod 3, say *N* = 3*K*, the best option is to set two of the three values to 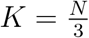 and make the remaining one equal to 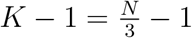. In summary, the maximum is attained when |*n*_0_ − *n*_1_| ≤ 1, |*n*_0_ − *n*_2_| ≤ 1, |*n*_1_ − *n*_2_| ≤ 1. We call a node which satisfies this condition a **tricenter** of the tree. Not all trees have one - for instance, the complete binary tree on *n* = 4 tips does not have one. The uniqueness of a tricenter follows from the uniqueness of the centroid, proven as part of the next result. Furthermore, the value of betweenness centrality at a tricenter is close to 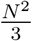, while having *n*_0_ = 0 (as would be the case for the root) results in a value of at most 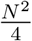, which is clearly suboptimal. Lastly, it is also clear that *n*_1_ = *n*_2_ = 0 (as would be the case for any tip) results in a betweenness centrality value of 0, the minimum possible.
b. Let *u* be the node of a phylogenetic tree *T*_0_ that has the smallest farness; we will show that *u* must be a centroid, and conclude by showing that every phylogenetic tree has a unique centroid. For the first part, let us consider the tree *T* (which will not be binary if *u* is not the root) obtained by rerooting *T*_0_ at *u*. Note that the distances in *T* are equal to the distances in *T*_0_. To make things general we consider a tree with branch lengths (weights). Let *v* be any child of *u* in *T*. Note that *d*(*v, x*) = *d*(*u, x*) − *w*(*uv*) for any node *x* in the subtree *T_v_* rooted at *v*, and *d*(*v, x*) = *d*(*u, x*) + *w*(*uv*) for any node *x* outside *T_v_* (including *u* itself). If we denote by *F* (*x*) the farness of a node *x*, then

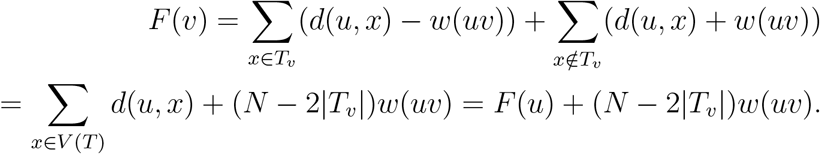 This recurrence is in fact the basis for a linear-time algorithm for computing *F* (*x*) for all nodes *x* in *T*_0_, starting from the root. Recall that *w*(*uv*) > 0 by assumption. Therefore, in order for *F* (*u*) to be the smallest farness, we must have

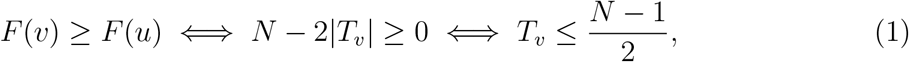

since *N* = 2*n* − 1 is odd, and in fact, the inequalities must be strict. It follows that the subtree *T_v_* of *T* rooted at any child *v* of *u* must contain fewer than half of all the nodes; therefore, *u* must satisfy the definition of a centroid. We note that two closely related results appear in the work on 1-medians of tree networks [21]. It remains to show that the centroid exists and is unique in a phylogenetic tree. For the existence, we apply a result by Camille Jordan [29] that proves the existence of a *separator node v* in any (not necessarily binary) tree *T*, which is a node defined by the property that any connected component obtained by removing *v* from *T* has size at most 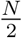, where *N* is the number of nodes in *T*. The reason that such a separator node is a centroid is that the connected components resulting from removing *v* are identical with the subtrees rooted at *v* if *T* is rerooted at *v*, and that *N* is odd, so 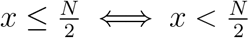 for *x* ∈ ℕ. For the uniqueness (in phylogenetic trees), suppose that this there is another node in *T*_0_, say *w* ≠ *u*, that also has the defining property of a centroid. Consider the connected component *C^u^* of *T*_0_ after *w* is removed that contains *u*, and the connected component *C^w^* of *T*_0_ after *u* is removed that contains *w*. Since *u* and *w* are centroids, *C^u^* and *C^w^* each contain fewer than half the nodes, so their union does not contain some node *v* of *T*_0_. But this cannot happen because if *d*(*u, v*) ≤ *d*(*w, v*) then *v* ∈ *C^u^*, while if *d*(*w, v*) ≤ *d*(*u, v*) then *v* ∈ *C^w^*. Lastly, we note that for any internal node *v*, there is some tip *w* such that the path from *v* to *w* strictly increases in farness; indeed, it is sufficient to always pick the child of the current node whose subtree contains fewer than half of all the nodes, by equation (1). Therefore, the maximum farness is attained at a tip, not at an internal node. To connect the concepts of centroid and tricenter, we note that a tricenter, if one exists, is also a centroid, by definition; in particular, a tricenter, if it exists, is always unique; however, a phylogenetic tree always has a centroid, while it may not have a tricenter.
c. Recall the assumption that the tree *T* has arbitrary positive branch lengths. Let *v* be a tip with a parent *u*. The defining equations for the Perron-Frobenius eigenvector specialized to the entries corresponding to *u* and *v* say that

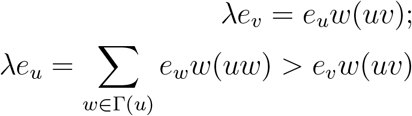

where the inequality follows from the fact that *u* has at least one other neighbor, and all the branch lengths and entries of 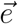 are positive. By cross-multiplying the inequality with the equality, we get

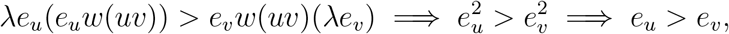

since *λ* > 0 and *w*(*uv*) > 0. Hence, a tip cannot contain the maximum eigenvector centrality, which is therefore attained at an internal node.

**Figure S6:**
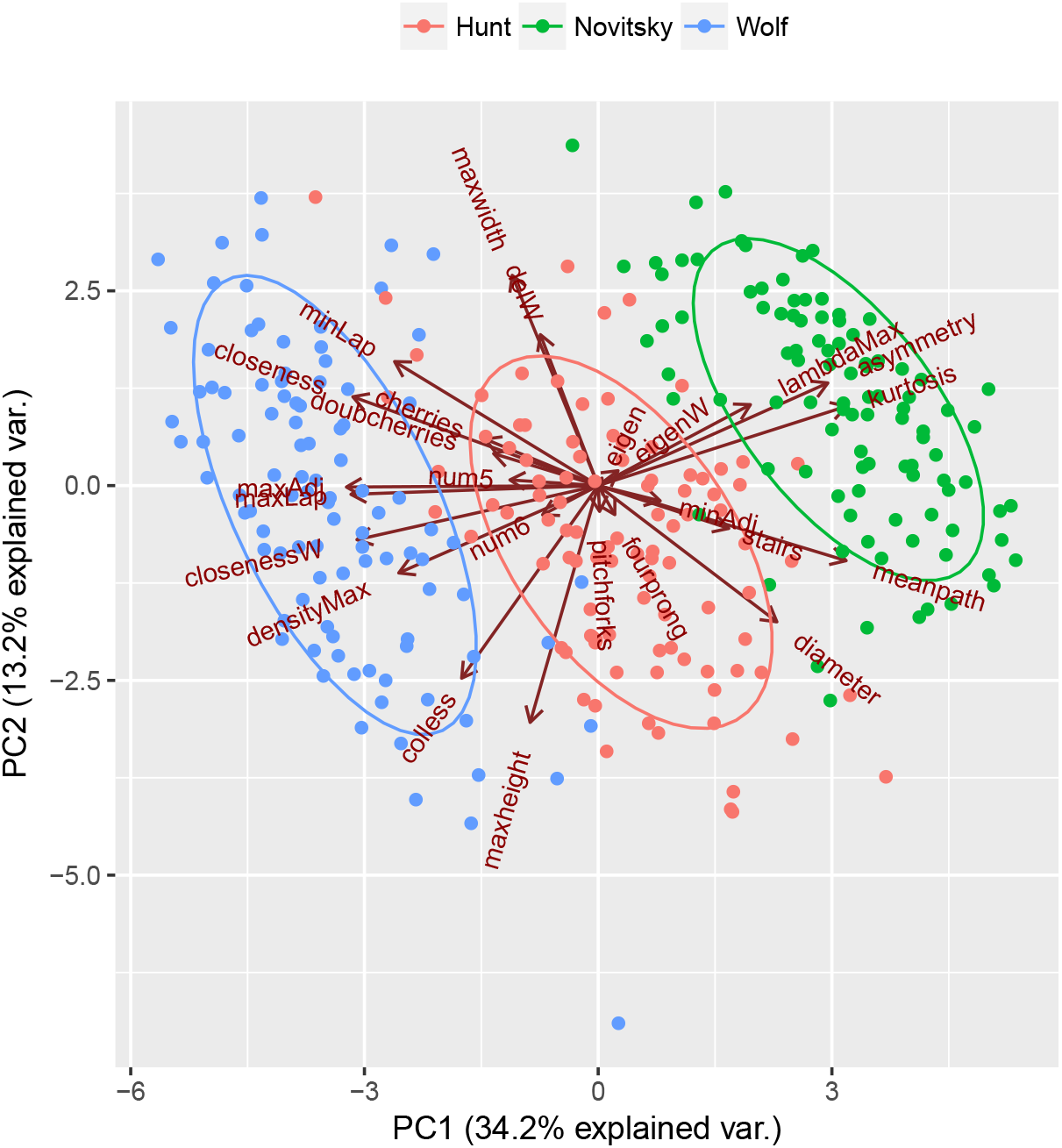
PCA biplots illustrating the separation among HIV phylogenies sampled from by using tree shape statistics.

**Figure S7:**
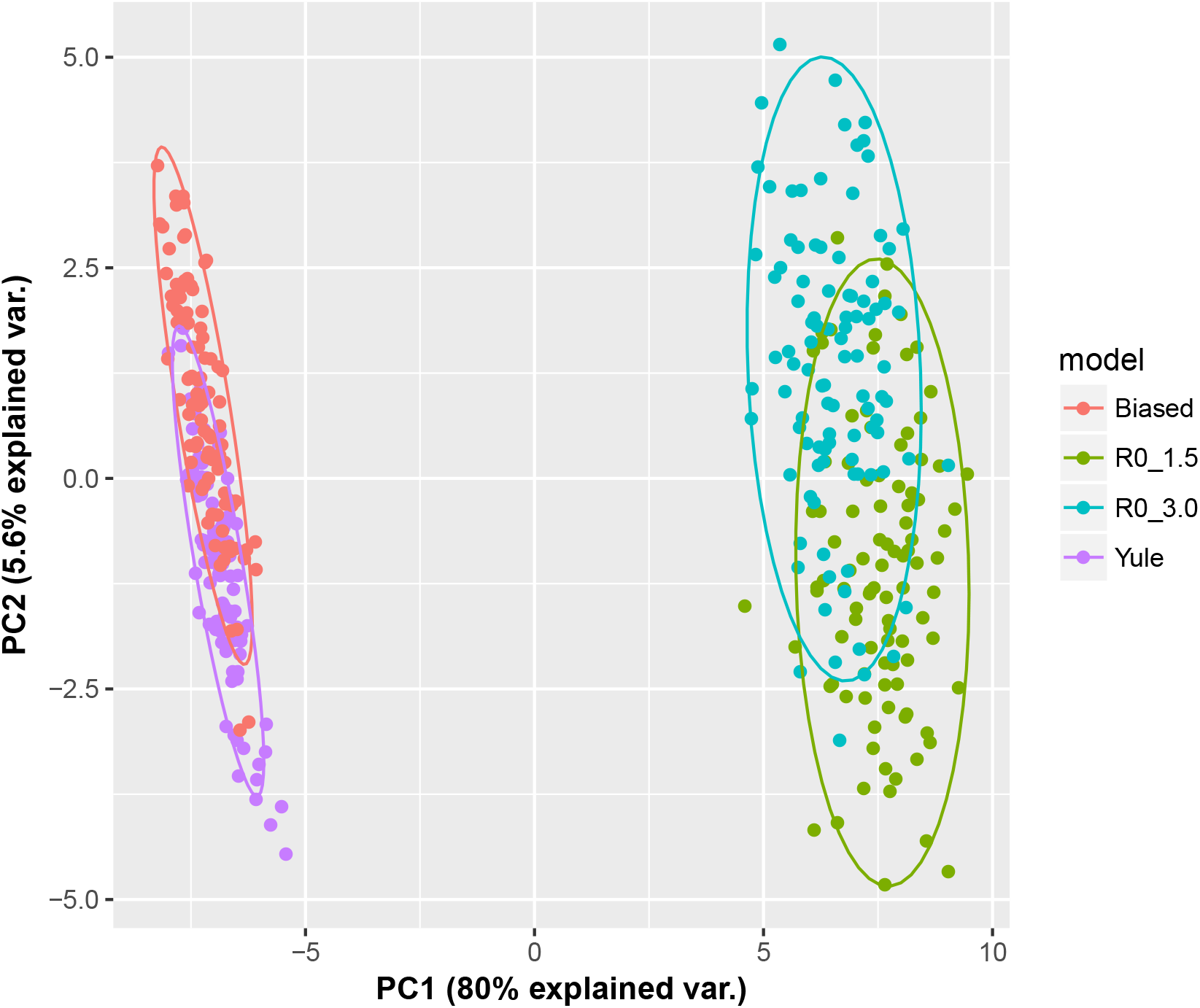
Kernel PCA plot illustrating the separation among phylogenies simulated from a biased model (red), a Yule model (purple) and two birth-death models (green and blue) by using the kernel method.

**Figure S8:**
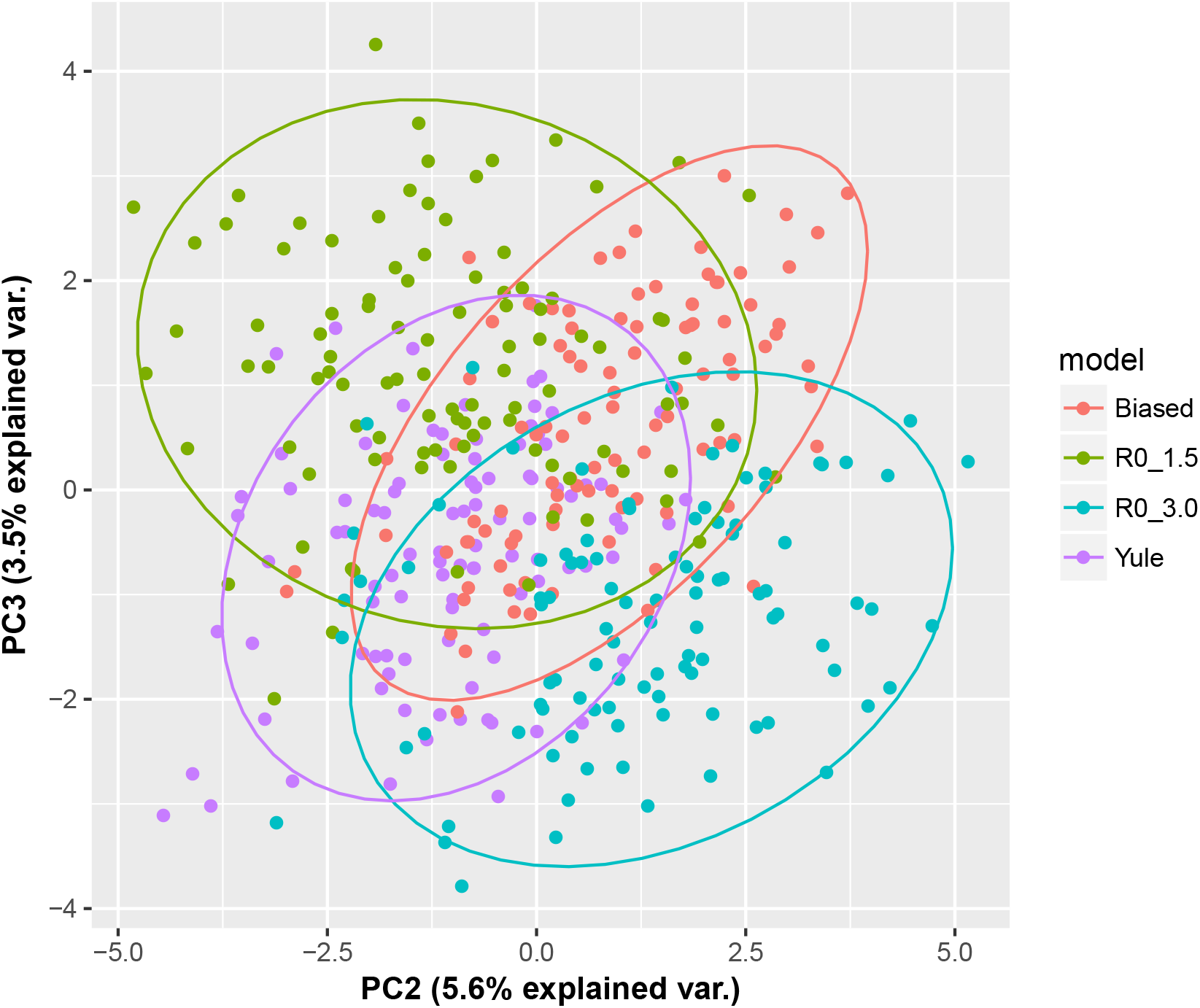
The same kernel PCA plot as above, components 2 and 3.

### 6.8 The distance Laplacian spectrum does not uniquely define a tree

We give an example of a pair of trees on *n* = 3 tips, with different branch lengths, whose distance Laplacian matrices are co-spectral. We obtained them by exploring all possible integer branch lengths, in order of total branch length (from smallest to largest), and used Maple [46] to check, for each resulting weighted topology, the existence of a different set of branch lengths that would result in a distance Laplacian matrix having the exact same characteristic polynomial, and hence the exact same spectrum. The first tree is therefore the tree with integer branch lengths having the smallest possible total length and admitting a co-spectral tree, the second tree, with respect to the distance Laplacian matrix.

**Figure S9:**
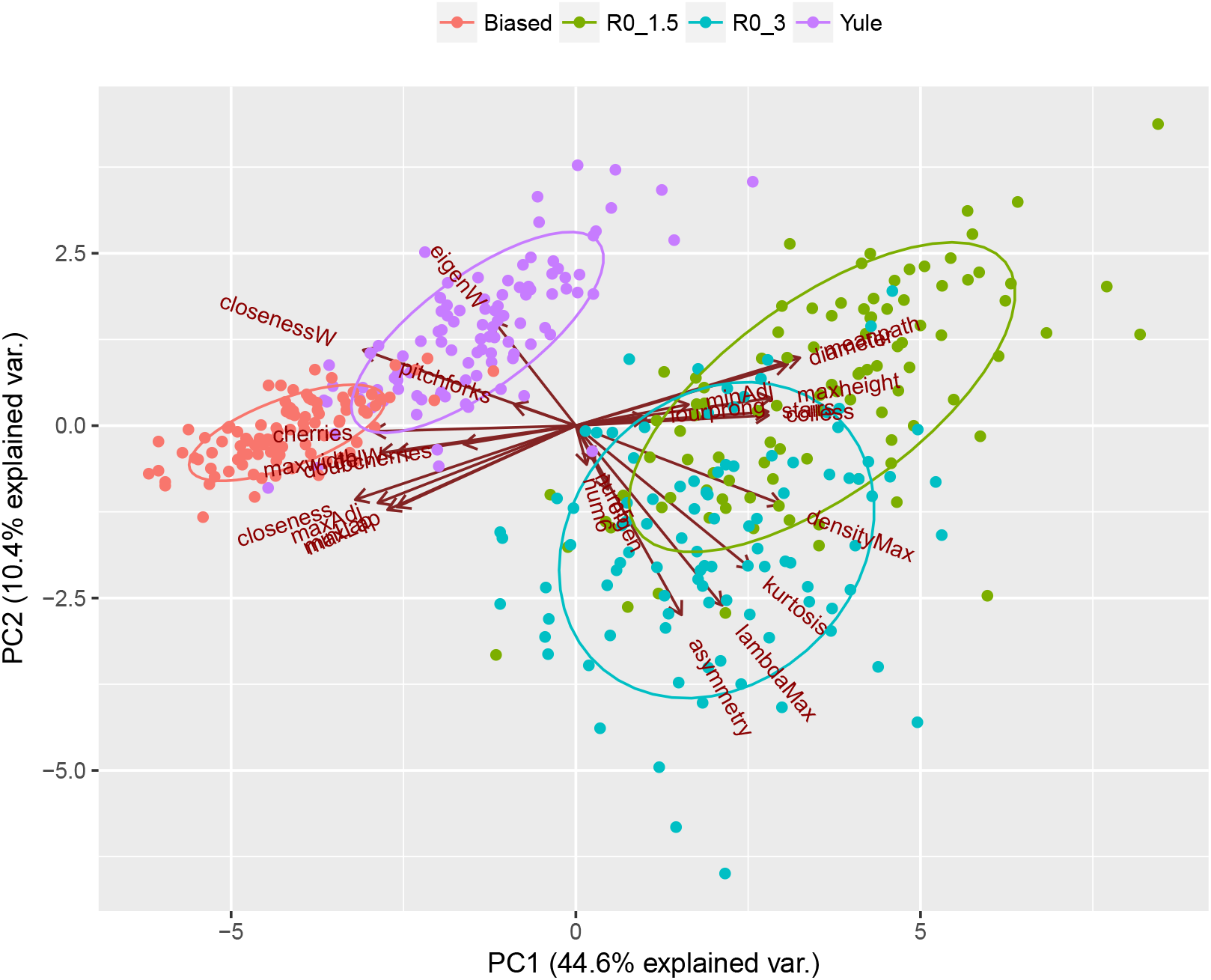
PCA biplots illustrating the separation among phylogenies simulated from a biased model (red), a Yule model (purple) and two birth-death models (green and blue) by using tree shape statistics.

The first tree has branch lengths *a* = 2 from the root to the first tip, *b* = 2 from the root to the internal node, and *c* = *d* = 1 from the internal node to the remaining two tips. The second tree has corresponding branch lengths *a* ≈ 0.16754417767858494454487, *b* ≈ 3.5136399160176469472222, *c* ≈ 1.3164466748555757609955, *d* ≈ 0.2455492734393688736927 (the exact expressions involve the roots of a polynomial of degree 22 and are provided in the Maple worksheet as a **Supplementary File**). These results were verified using the *RPANDA* package [48] in R.

### 6.9 Distance Laplacian spectra do determine small unweighted trees

We created a computer program to exhaustively test whether any pair of distinct phylogenetic trees had identical distance Laplacian spectra. The trees were generated by the method proposed by Gang Li in his Masters thesis [37], which only requires enough space for a single tree and a constant amount of operations per tree generated. For each tree we computed the distance Laplacian spectrum and compared it to that of all the other trees of the same size. Since the number of unlabelled phylogenetic trees is known to grow as *Cα^n^n*^−3/2^, where *C* ≈ 0.792 and *α* ≈ 2.483 [37], it quickly becomes infeasible to enumerate and keep in memory the full spectra of the distance Laplacian matrix for each possible tree. For this reason we made use of several optimizations.

First, we observed that two trees can only have cospectral distance Laplacian matrices if the traces of these matrices, known as the Wiener indices of the trees (the sum of all node-node distances) [45], are equal. In particular, this means that the computation could be parallelized if the values of the possible Wiener indices were known in advance, together with their multiplicities. Second, we observed that the distribution of the Wiener indices of a tree on *n* tips could be pre-computed efficiently from the heights and Wiener indices of the subtrees composing it. Lastly, we were able to augment the tree generation algorithm with an additional set of instructions that allowed us to also compute the Wiener index while retaining the constant amortized time complexity.

**Figure S10:**
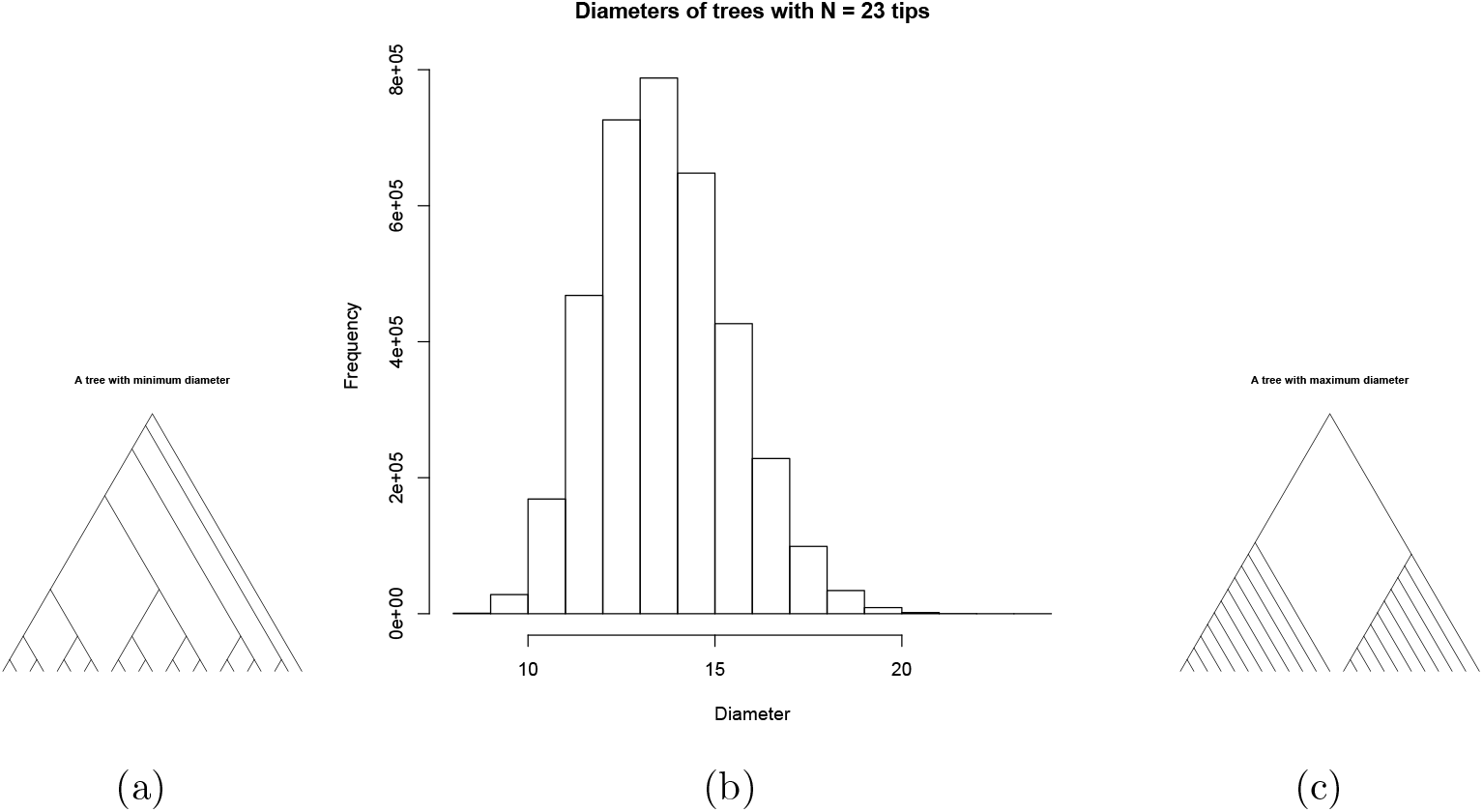
The distribution of diameters for phylogenetic trees on *n* = 23 tips, with two extremal trees

In this manner, we pre-computed the distribution of the Wiener indices, decided on a split of the trees between processes, and limited the spectral comparisons to those within the same process. We performed the comparisons themselves in two passes; in the first pass, all trees with appropriate Wiener index were generated and their highest and lowest eigenvalue stored in a large matrix; in the second pass, only those trees that had at least one match were re-generated, and their full spectra compared. The computation for *n* = 32 tips, involving 7.86 billion trees split between 10 processors, required approximately 2 weeks of computation on a cluster, and did not produce any co-spectral trees, although some of the spectra were within 0.01 distance in the max norm (*L*_∞_).

**Figure S11:**
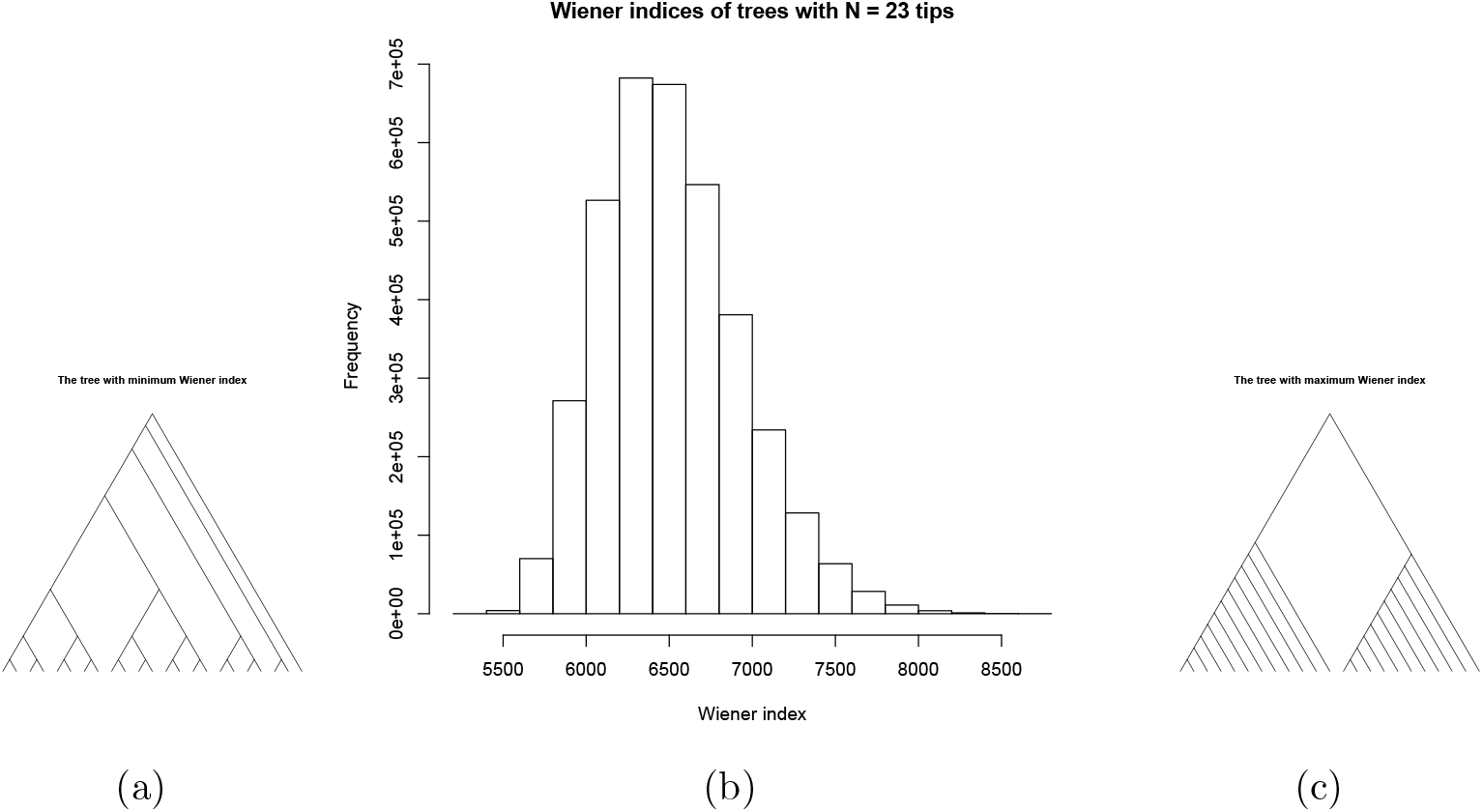
The distribution of Wiener indices for phylogenetic trees on *n* = 23 tips, with two extremal trees

**Figure S12:**
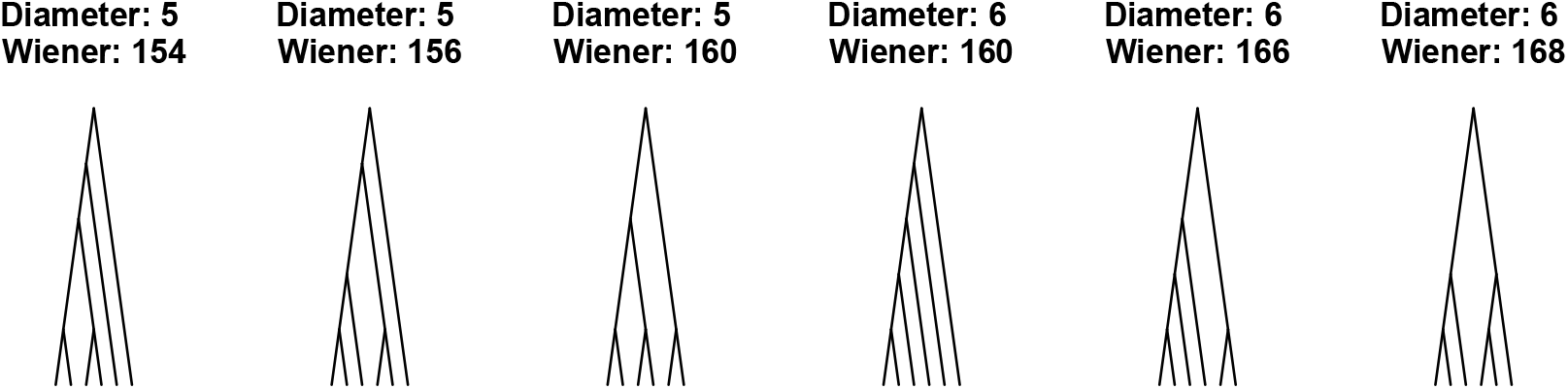
The 6 phylogenetic trees on *n* = 6 tips, with their diameter and Wiener index

**Figure S13:**
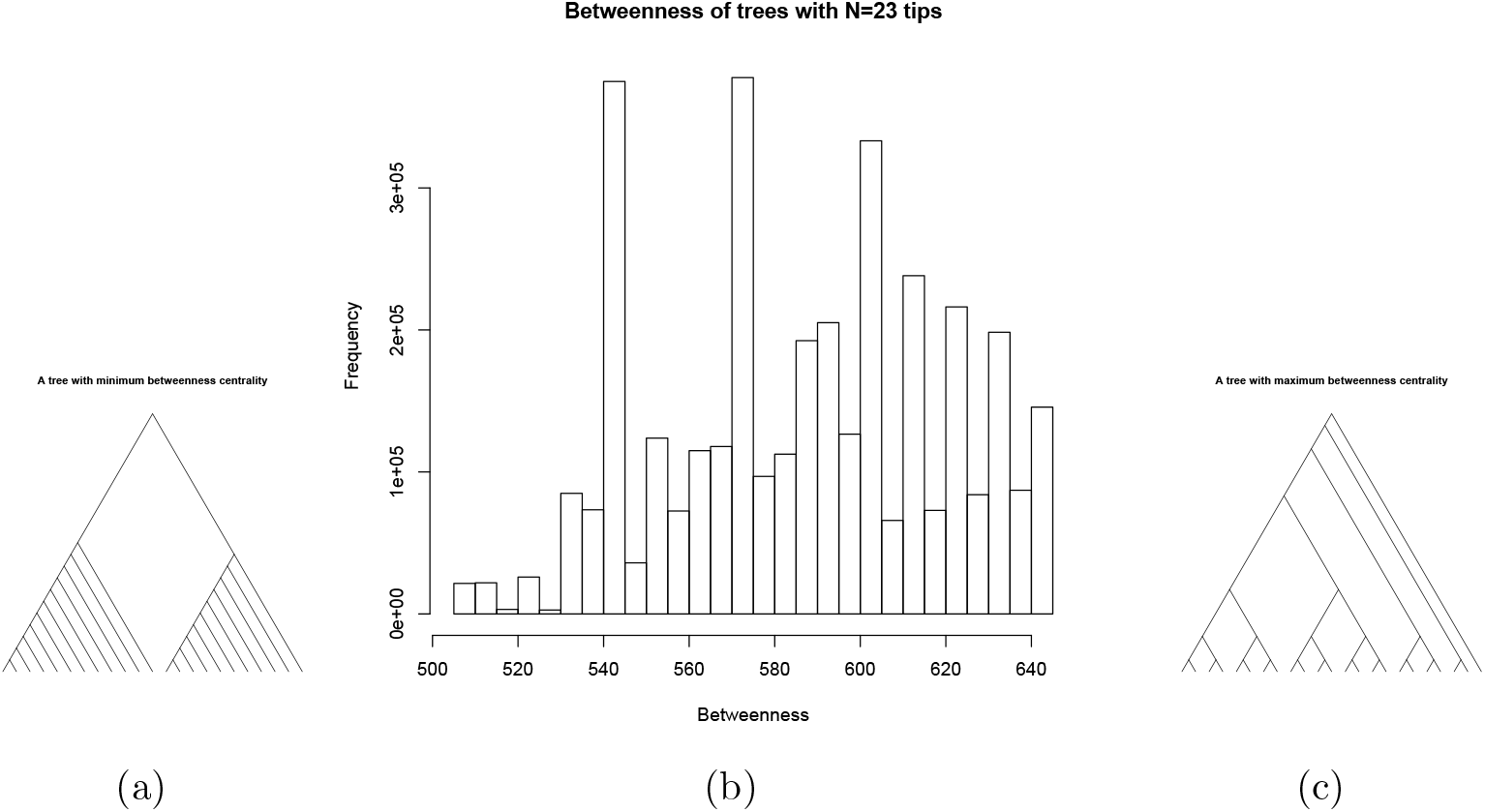
The distribution of maximum betweenness centralities for phylogenetic trees on *n* = 23 tips, with two extremal trees

**Figure S14:**
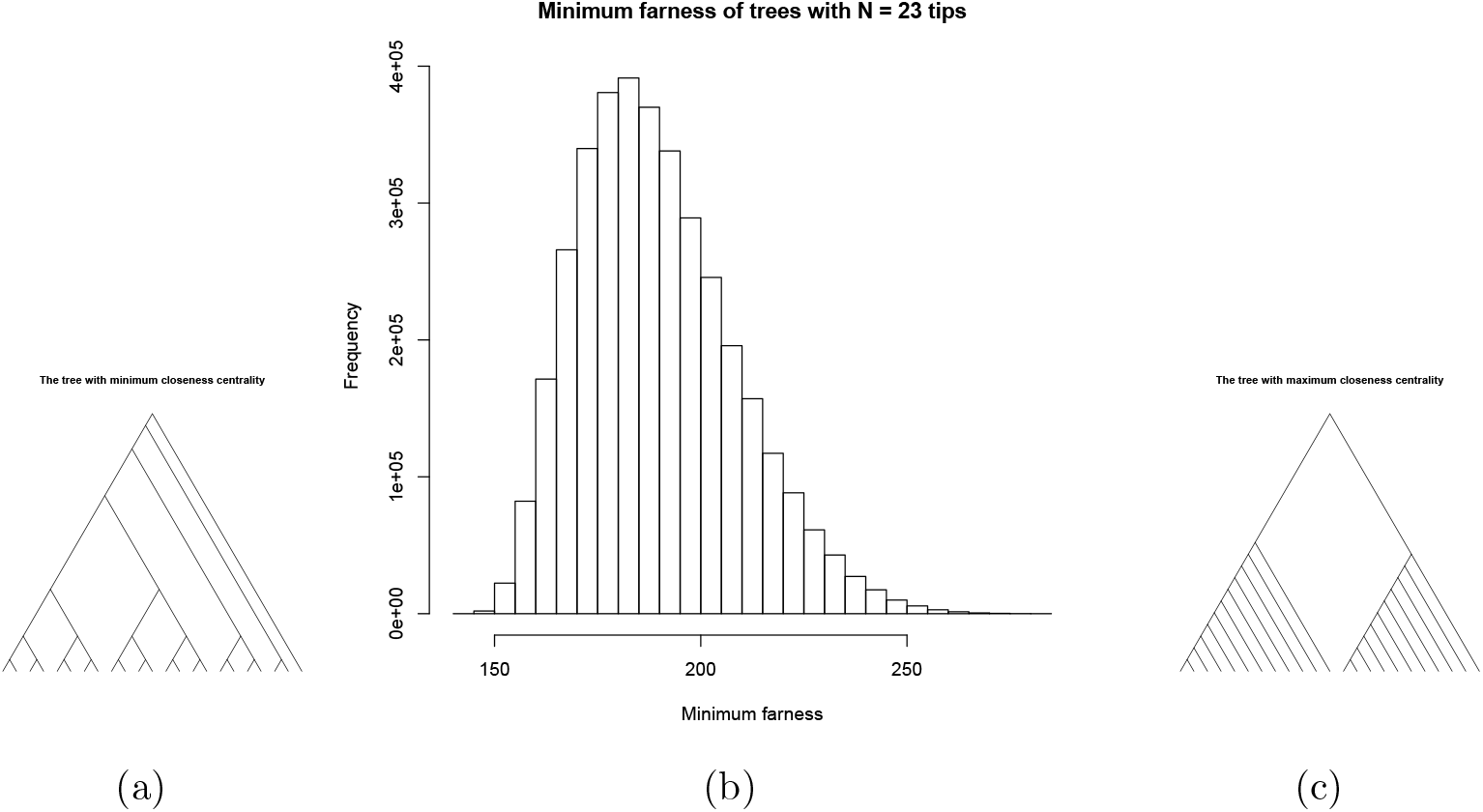
The distribution of minimum farness (inverse of maximum closeness centralities) for phylogenetic trees on *n* = 23 tips, with two extremal trees

**Figure S15:**
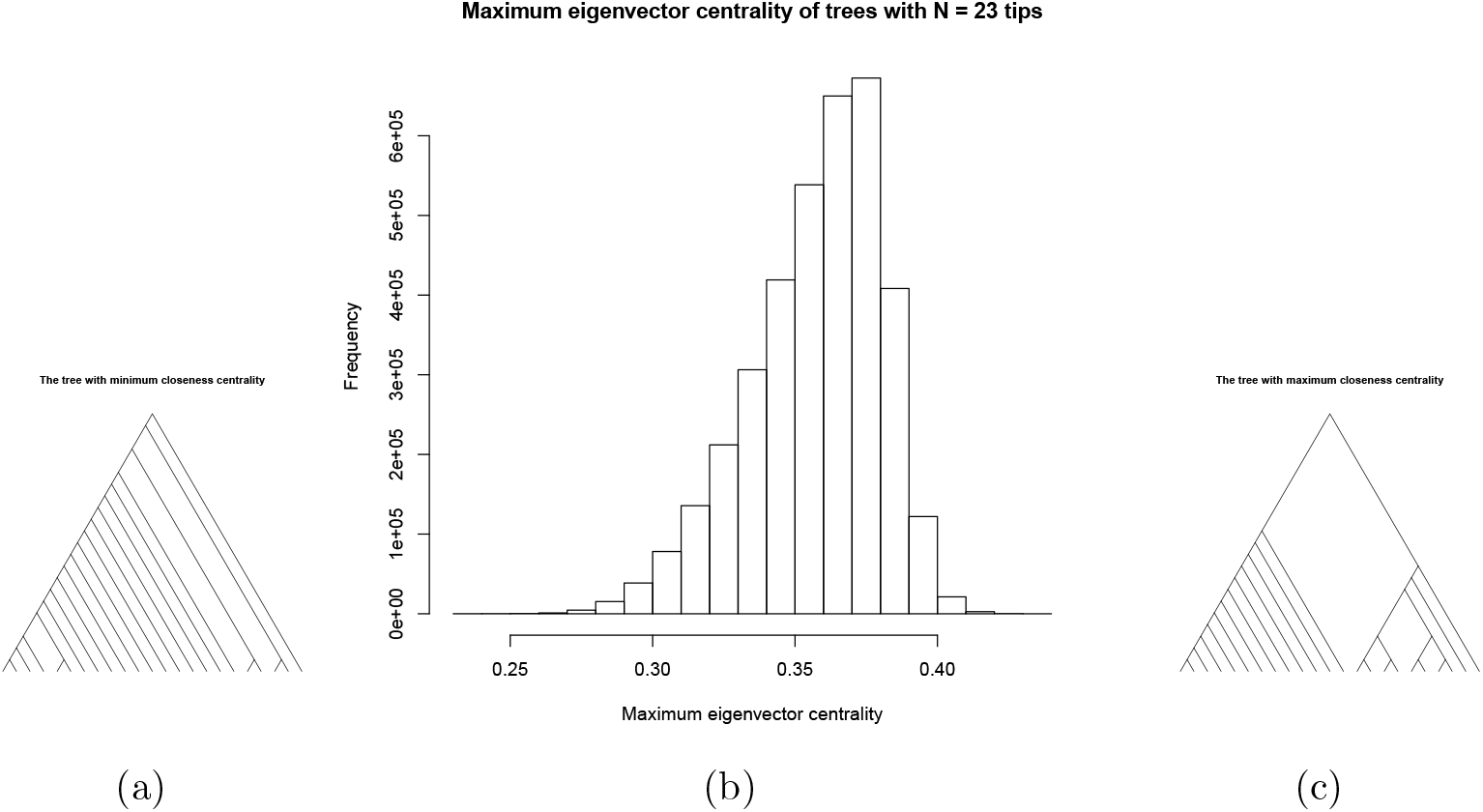
The distribution of maximum eigenvector centralities for phylogenetic trees on *n* = 23 tips, with two extremal trees

**Figure S16:**
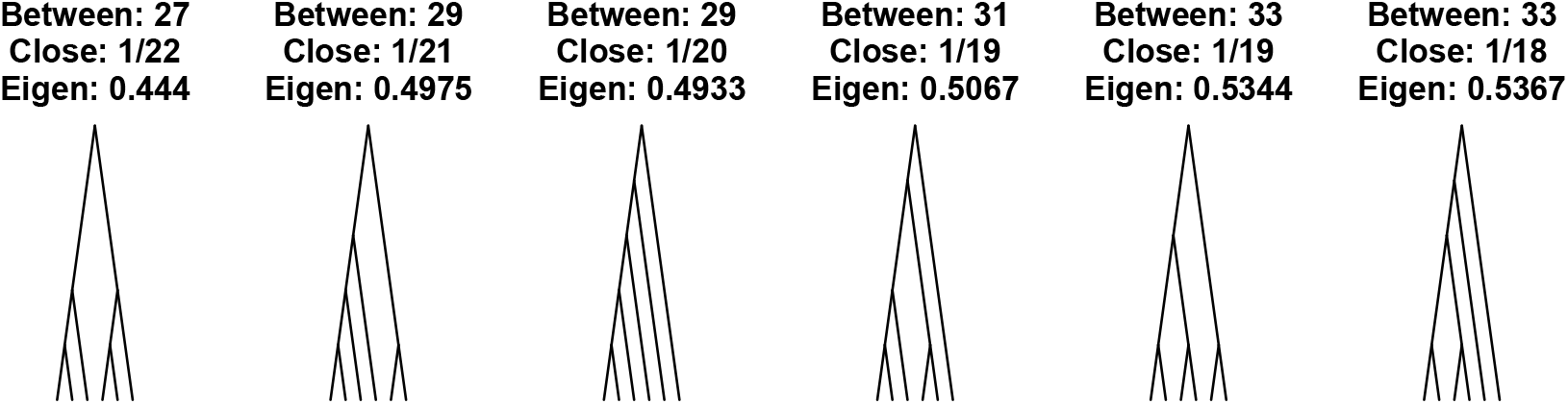
The 6 phylogenetic trees on *n* = 6 tips, with their betweenness, closeness, and eigenvector centrality

## Notes

http://github.com/Leonardini/treeCentrality

http://github.com/Leonardini/treeCentralityData

### Competing Interest Statement

The authors have declared no competing interest.

### Summary of Updates

- Additional experiments to identify feature importance - A more unified evaluation of tree shape statistics - Additional evaluations on various simulation models

